# “What’s SUPP” developing an *in vitro* model for healthy oral biofilms

**DOI:** 10.64898/2026.06.19.733444

**Authors:** A. Labossiere, M. Ramsey

**Author notes:** <u>Author Contributions:</u> MR and AL designed the research, AL performed the research. AL and MR wrote the paper. <u>Competing Interest Statement:</u> There are no competing interests.

## Abstract

Human supragingival plaque (SUPP) is a polymicrobial biofilm whose contents undergo dysbiotic transitions during multiple oral diseases. The study of healthy SUPP may lead to future pro or prebiotic therapies, to help prevent or revert dysbiosis during disease. However, many oral plaque models focus on the cultivation of oral pathogens and do not well cultivate commensal SUPP populations. Here, we use a 16S microbiome guided iterative approach to develop a low-cost high sample number SUPP model. Our model demonstrates several findings including a surprisingly minimal impact on salivary preparation methods on model microbiota and the ability to test microbial interactions with added oral strains to assess their fitness. This model provides a reductionist system for the study of healthy oral commensals in a complex polymicrobial framework in the absence of host immune responses.

## Introduction

The field of microbiology has largely, and necessarily, been developed with a focus on infectious disease investigation and prevention. Recent efforts in microbiome research have described healthy commensal microbial communities (Huttenhower et al. 2012) yet their role and interactions (microbe-microbe / microbe-host) in health are poorly understood. Opportunistic pathogens involved in oral diseases such as periodontitis and caries are present in healthy individuals typically at low abundance (Bertolini et al. 2022; Bowen et al. 2018). While the transition from health to oral disease is strongly influenced by host behaviors and systemic disease(s), there is likely a role of the healthy commensal microbiota that can be leveraged to prevent or minimize oral disease. This is supported by cases where pathogen carriage may be elevated in some individuals where disease remains absent (e.g. high *S. mutans* carriage in caries free individuals, and in the fortunate failure for many to transition from gingivitis to periodontitis) (Zhang et al. 2022; William G. Wade 2021).

The cooperative nature of oral pathogens during disease is well understood (Kwon et al. 2021) (Rathee and Sapra 2025; Spatafora et al. 2024; Caufield et al. 2015) as is the transition of healthy oral microbial communities to those observed in disease (Loesche et al. 1977; George Hajishengallis et al. 2012; G. Hajishengallis and Lamont 2012; P. D. Marsh 1994). We and others have also observed the influence of commensal microbes on the behavior of pathogens, both oral and extra-oral, partially explaining the presence of pathogens in healthy microbiomes (Ramsey and Whiteley 2009; Philip D. Marsh 2006; Eren et al. 2014). Mechanistic studies of interaction are largely reductionist, and while they provide valuable data, are limited especially in the context of oral <u>sup</u>ragingival <u>p</u>laque (SUPP), which are microbially diverse, highly structured biofilms residing on the tooth surface (Jessica Mark Welch 2022; Eren et al. 2014). There is a need to extend initial reductionist mechanistic findings between individual species into more comprehensive models to determine if *in vitro* interaction mechanisms are meaningfully relevant in the context of their greater microbiome community. *In vivo* models in humans are costly and limited in what experimental interventions can be utilized. Animal models possess the same shortcomings and are complicated by unique host properties as well as their native microbiota.

There are a plethora of *in vitro* SUPP models utilized to date (Pigman et al. 1952; Russell and Coulter, 1974; Keevil et al. 1987; Bradshaw et al. 1996; Guggenheim et al. 2001; Edlund et al. 2013; Naginyte et al. 2019). While these are suitable for the study of oral pathogens, we assert here that they are insufficient for the study of healthy SUPP communities. One drawback is that many of these models utilize anaerobic culture conditions, heavily selecting for subgingival species and enrichment of periodontal pathogens (Dini et al. 2023; Adami et al. 2025; Ghesquière et al. 2023; Bao et al. 2015; Wang et al. 2024). Other drawbacks include using growth media that do not recapitulate a saliva-rich SUPP environment (Naginyte et al. 2019; Edlund et al. 2013), are pre-assembled communities of species present in SUPP that are not highly abundant in health, or do not use SUPP as an inoculum (M. Guggenheim et al. 2001; Guggenheim et al. 2004; Giertsen et al. 2011; Shapiro et al. 2002). In cases where microbiome observations of these models are available, they do not appear to contain bacterial species in relative abundances that are similar to that of healthy SUPP (B. Guggenheim et al. 2001; Deari et al. 2025; Tsutsumi et al. 2018). This led us to develop an *in vitro* SUPP model specifically for the study of healthy commensal interactions.

Our criteria for an improved *in vitro* SUPP model was based on multiple factors; it should use environment and medium conditions reflective of health, utilize SUPP as an inoculum, incorporate host substrates present in SUPP, and allow for sample numbers required for well powered microbiome assessment. Here, we describe a low-cost, accessible, high sample number *in vitro* SUPP model. We used saliva coated hydroxyapatite discs incubated in a 24-well plate format under a thin layer of saliva-based growth medium, incubated aerobically under rocking motion. We designed this model based on iterative microbiome-based assessment, comparing initial SUPP donor microbiomes to those in differing medium and incubation conditions over 3-5 days to refine the model. We then used the refined model to compare the impact of saliva preparation methods on the model microbiome. We also incorporated fluorescent labeled bacterial strains into the SUPP community, allowing us to quantify the fitness of previously developed mutant vs wild-type strains to assess their relevance in a broader SUPP community. This model will be useful for researchers to study the impact of microbe-microbe interactions in a complex community independent of host immune responses and can allow for the investigation of both microbial mechanistic interactions as well as the influence on host saliva on changes in community composition.

## Results

### A saliva-based medium for SUPP cultivation

We began with a saliva-based medium that would permit growth of health-associated supragingival plaque (SUPP) organisms called “CORM” (complex oral rich medium). This consisted of commercial defined medium (Teknova EZ-Rich) amended with 25% sterile saliva, and added nucleotides (see materials and methods). CORM allowed robust overgrowth of underrepresented taxa present in our SUPP inoculum (data not shown). This led us to discover that SUPP species were amply growing on nucleotide and amino acids present in CORM and not saliva (Fig. 1). As we desired to utilize saliva as the primary source of carbon and energy for our SUPP community, we tested several media formulations to create an original saliva-supplemented complex medium (Fig. 1A-B). We compared the growth of SUPP in MOPS supplemented medium versus MOPS buffer alone. With 25% saliva as the sole carbon source, MOPS supplemented medium showed enhanced SUPP growth vs MOPS buffer alone (p=0.05), indicating it supported favorable growth.

**Figure 1.**
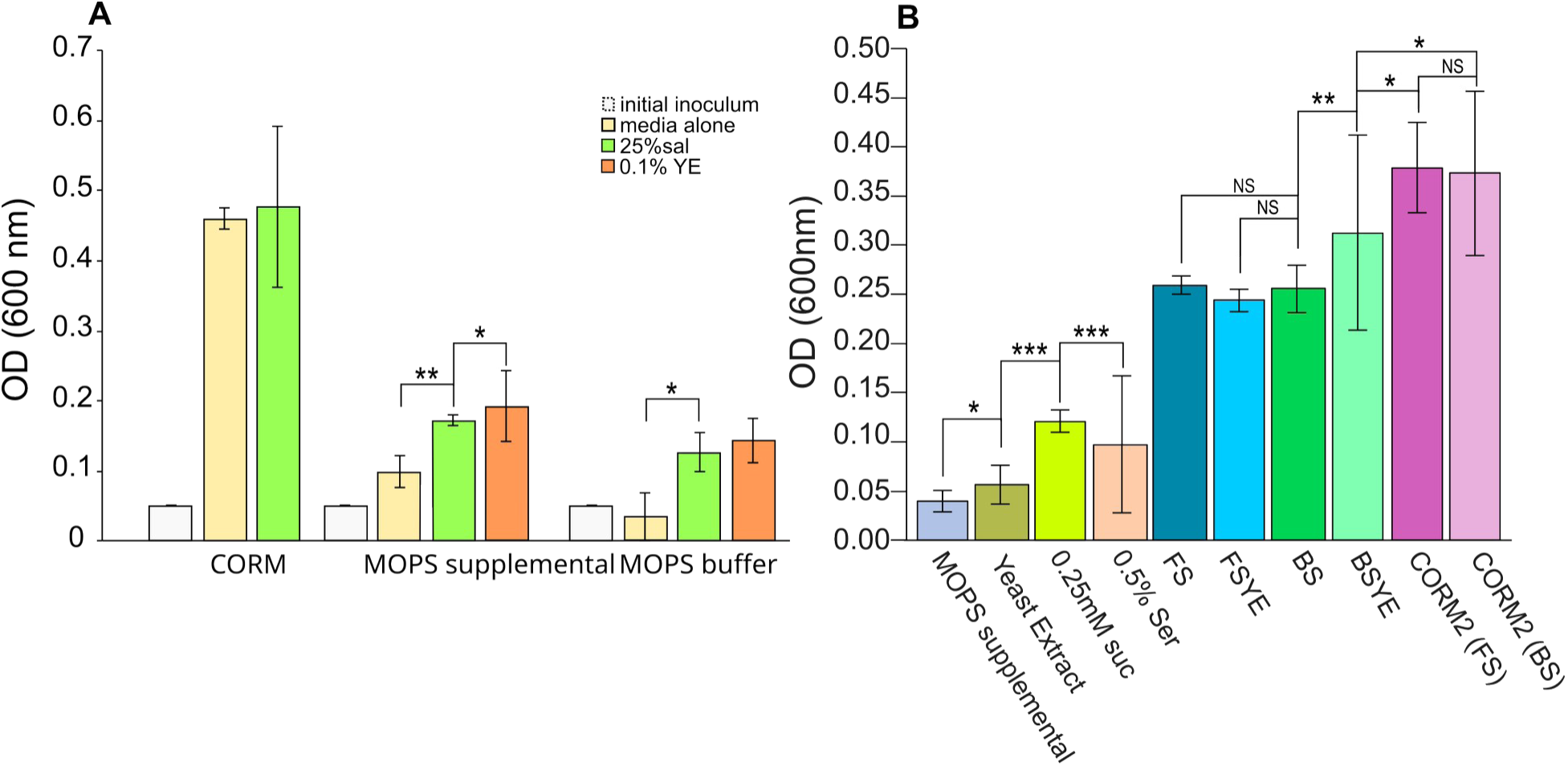
A saliva-based medium for SUPP cultivation. (A) Optical density comparison of CORM media, MOPS supplemented media and MOPS alone with: no additional carbon source, 25% saliva as a carbon source, and with further amendments (0.1% yeast extract, YE) to ensure saliva utilization within media formulation and to supplement potential auxotrophies. (B) Comparison of SUPP growth at 24h with different amendments along and comparison of boiled (BS) vs filtered saliva (FS). Significance determined by Student’s T-test, (n=3) *p ≤ 0.05, **p ≤ 0.01, and ***p ≤ 0.001.

We further amended our supplemented MOPS medium with other sources of carbon and energy available to SUPP bacteria (Fig. 1B). These included a low concentration of dietary sugar (suc – 0.25 mM sucrose), 0.5% heat inactivated human serum (ser) as a proxy for gingival crevicular fluid, and yeast extract (0.1% w/v). Yeast extract concentrations were low enough to supplement most cofactor auxotrophies but not enough to be utilized as a source of carbon and energy alone (Fig. 1B). We observed greater overall growth with these amendments added to a supplemented MOPS based medium with 25% saliva than without; and deemed this new formulation “CORM2”. Using CORM2 we were now able to examine if salivary preparation methods could have an impact on SUPP growth and composition *in vitro*.

### Saliva sterilization method has little effect on microbiome composition

Our initial *in vitro* SUPP model used CORM2 and healthy donor SUPP as an inoculum (see materials and methods under “model set-up” for full description). Briefly, we utilized saliva-coated hydroxyapatite (HA) discs covered by ∼1mm of CORM2 (350 μl) in each well of a 24-well plate and incubated under rocking motion in a 5% CO2 atmosphere at 37°C. SUPP was harvested from healthy adults that abstained from tooth brushing for 24hr prior to collection. Diluted SUPP from an individual donor was added to CORM2, incubated as described and HA discs were transferred to fresh medium every 24hr. We performed 16S microbiome characterization for donor SUPP and HA disc biofilm cultures at indicated timepoints. We used V1-V2 16S sequencing followed by QIIME2 analysis similar to our previous methods (Perera et al. 2022) further described in materials and methods and in supplemental data. Sample numbers were chosen based on power analysis calculations to ensure adequate sample size for variation based on preliminary sequencing from our initial model conditions (see supplemental for power calculations).

We then used our model to investigate how common saliva sterilization methods (vacuum filtering vs boiling) impacted SUPP community composition. We compared microbiome profiles for each media type (n=11) after 5 days (Fig. 2A) using vacuum filter sterilized vs boiled saliva at 25% within CORM2 in parallel from the same SUPP inoculum. Disappointingly, we observed major outgrowth of *Neisseria* species (>40% total abundance) in all tested conditions (Fig. 2A, “unfiltered sequences”). Many of our datasets contain outgrowth of organisms such as *Neisseria*, *Enterobacteriaceae*, and *Lautropia mirabilis*. Fortunately, we observed considerably less outgrowth at 3 days in subsequent experiments, and also found that in prior human *in vivo* dental material implant experiments had considerable growth of *Neisseria* (Tu et al. 2020; Utomo et al. 2024). Thus, while imperfect, our model showed similar results to oral substrates used in human tests *in vivo*.

**Figure 2.**
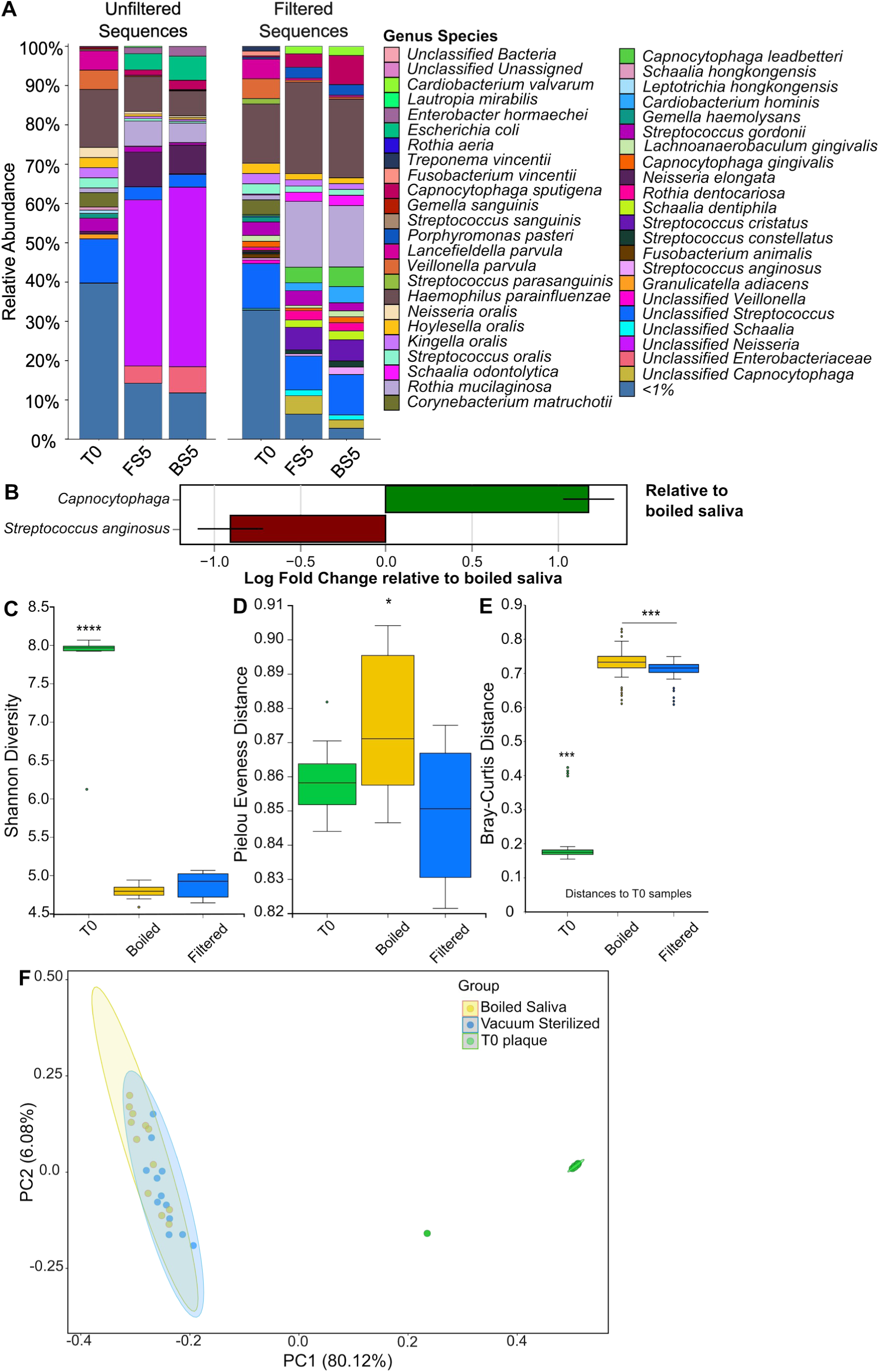
Minor community impact by salivary preparation method. (A) DADA2 species assigned 16S profiles of the original donor SUPP (T0), 5 day model cultures with filter-sterilized saliva (FS5) or boiled-sterilized saliva (BS5) were compared. Highly abundant *Neisseria* sequences were computationally removed (Filtered Sequences) to reveal highly similar, diverse underlying microbiota. All following figures depict filtered-sequence data, unfiltered data comparisons and remaining metrics are in Figs. S1, S10. (B) ANCOM-BC comparisons between FS5-BS5 reveal only two significantly differed taxons for either preparation method. (C) Shannon diversity and Pielou’s Evenness (D) indicate broad similarity between either saliva preparation method in the model microbiome as do β-diversity metrics for Bray-Curtis (E) and unweighted Unifrac comparisons (F). α and β-diversity metrics utilized Kruskal-Wallis one-way ANOVA and pairwise PERMANOVA analyses respectively. n=11 for all comparisons, *p ≤ 0.05, **p ≤ 0.01, and, ***p ≤ 0.001. Ellipses indicate a 99% confidence interval.

Filtering *Neisseria*-aligned sequences from this data allowed further resolution of the underlying community composition (Fig. 2A “filtered sequences”). In both saliva conditions, SUPP taxa including *Streptococcus* species (41%), *Haemophilus parainfluenzae* (23%), *Rothia mucilaginosa* (15.5%), *Capnocytophaga* species (9%) *Porphyromonas pasteri* (2.72%), *Cardiobacterium valvarum* (2%), and lesser amounts of anaerobic organisms such as *Veillonella* (0.5%) constituted ∼94% of the biofilm population after read filtering. This demonstrated that the remaining microbial composition is highly similar to that found in the initial SUPP inoculum. We then compared each condition to detect differentially abundant organisms via <u>A</u>n A<u>n</u>alysis of <u>C</u>omposition <u>o</u>f <u>M</u>icrobiomes with <u>B</u>ias <u>C</u>orrection (ANCOM-BC) (Lin and Peddada 2020) (p = 0.0001), (Fig. 2B). Interestingly in filter-sterilized saliva samples, the genus *Capnocytophaga* was increased (1.2-log fold change) and a decrease of *Streptococcus anginosus* (0.9-log fold) was observed, showing otherwise insignificant differences in between saliva preparation methods on SUPP microbiome composition.

α-diversity comparisons showed that T0 SUPP inoculum had a Shannon index value of 7.8, indicating high richness and evenness, compared to the median value of boiled (4.78), and filtered saliva samples (4.87) (Fig. 2C), suggesting similar community composition for both. Boiled saliva samples had a slight decrease in Chao1 index values compared to filtered (p<0.0001) (Supplemental 1A), suggesting filtered saliva retains a slightly wider variety of rarer species while both communities contain relatively high species evenness (Fig. 2D). Filtered saliva samples contain less species evenness than our T0 inoculum differing by 2% on a Pielou Evenness index score (p=0.12) (Fig. 2D). As expected, our T0 sample contained the highest Faith’s PD score (Supplemental Fig. 1B), with filtered saliva samples higher than boiled (0.5-point difference) (p<0.005).

For β-diversity, Bray-Curtis measurements (Fig. 2E) indicated that the filtered and boiled saliva samples differed in community composition compared to inoculum and were more similar to each other, consistent with unweighted Unifrac comparisons (Fig. 2F). While both preparation methods had highly similar results, filtered saliva had slightly favorable comparisons to the initial inoculum and we opted to use filtered saliva for the remaining model conditions.

### CORM2 medium better approximates SUPP communities compared to SHI in vitro

We examined the influence of SHI medium, a complex blood-containing medium used to cultivate subgingival pathogens, on the composition of healthy SUPP in our model. Previously, SHI has been used to cultivate both supra- and subgingival plaque species (Du et al. 2021; Baraniya et al. 2020; B. Li et al. 2017; Edlund et al. 2013; Lamont et al. 2021; Tian et al. 2010), but the original paper (Tian et al. 2010) does not utilize it for SUPP cultivation, instead using saliva inoculum. Given its widespread use, we wished to compare its performance vs CORM2 in our model. Figure 3 details comparisons of *in vitro* model SUPP culture from an independent donor for SHI vs CORM2 media (additional subjects shown in supplemental S2-S4). Inter-individual variation of SUPP inoculum was diverse as expected and *Neisseria* overgrowth was again present. After filtering of *Neisseria* reads, the remaining community composition was again highly similar to each donors SUPP inoculum. Interestingly, each SUPP inoculum had *Lautropia mirabilis* at <1% total abundance, but the saliva-based medium samples contained this organism at 5-18% total abundance after 3 days of cultivation. Interestingly, two participants samples cultivated in SHI had overgrowth of *Aggregatibacter* species such as *A. kilianii* and *A. aphrophilus*. Overall, SHI medium samples contained a typically lower plaque diversity compared to CORM2 samples in the SUPP model.

**Figure 3.**
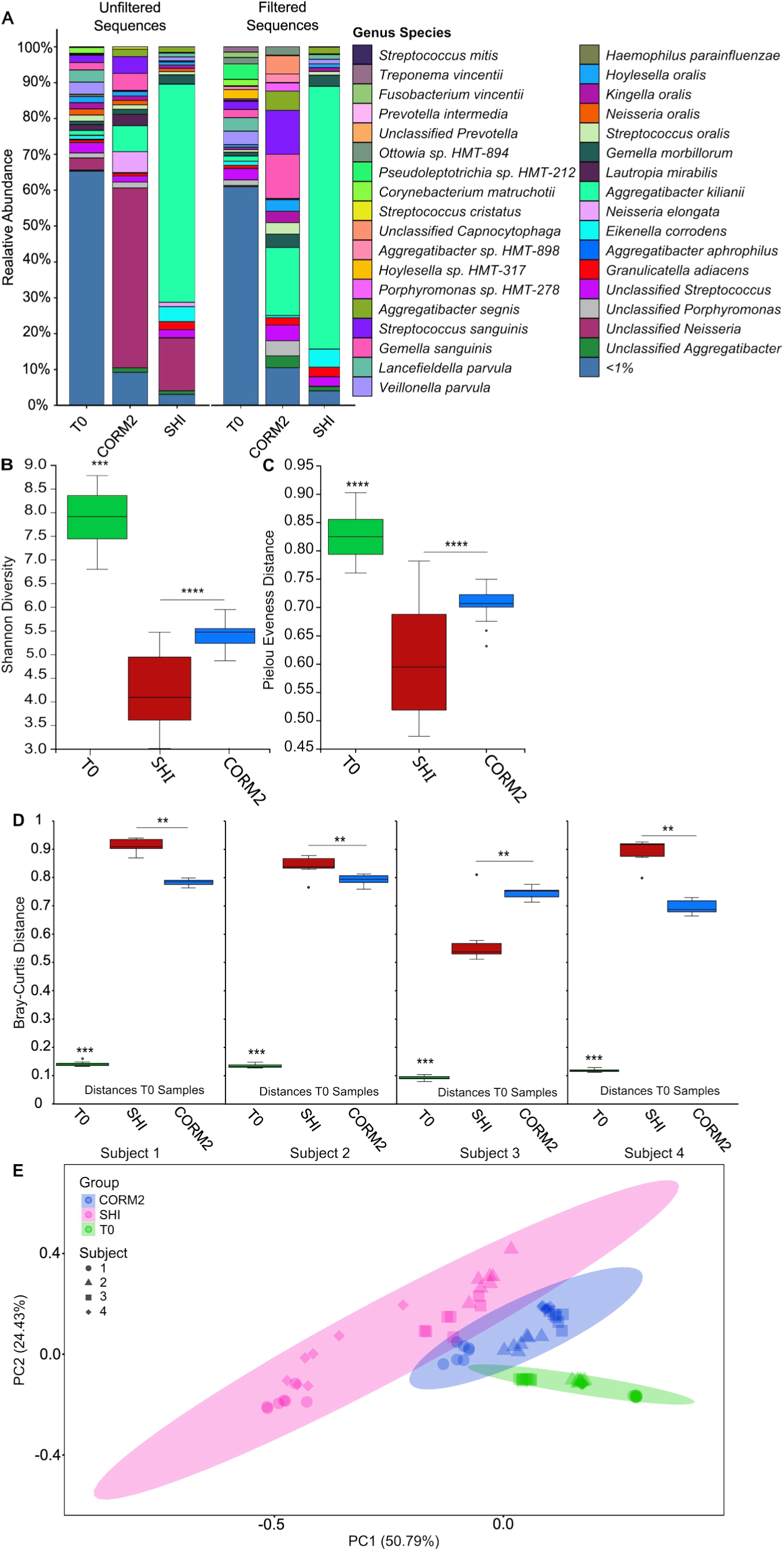
Medium type considerably impacts model microbiomes. (A) DADA2 species assigned 16S profiles of subject 1 donor SUPP (T0), compared to 5-day model growth in saliva-based CORM2 or Shi medium. Highly abundant *Neisseria* and *L. mirabilis* reads were computationally removed (Filtered Sequences) to reveal the underlying microbiota. Remaining subject data are in Figs. S2-4 and unfiltered diversity metrics comparisons are in Fig. S11. (B) Shannon diversity and Pielou’s Evenness (C) indicate both medium conditions differ from inoculum with CORM2 conditions more similar to T0. (D) Bray-Curtis data reveals similar patterns for all but Subject 3 and unweighted Unifrac comparisons indicate differing community structure for all 3 conditions across subjects with CORM2 samples more similar to T0. α and β-diversity metrics utilized Kruskal-Wallis one-way ANOVA and pairwise PERMANOVA analyses respectively. n=7 for each comparison for 4 separate subjects, *p ≤ 0.05, **p ≤ 0.01, *** p ≤ 0.001, and **** p ≤ 0.0001. Ellipses indicate a 99% confidence interval.

By examining diversity metrics in aggregate per medium rather than per individual we gain a deeper understanding of the community makeup. In Figure 3B, Shannon diversity of CORM2 cultures had less variability (p<0.001) while composition of phylogenetic diversity was more variable in weighted Unifrac analysis (Figure 3E). The greatest variability was observed in evenness for SHI medium (Fig. 3C). For β-diversity metrics comparing each subject to their respective T0 sample (Fig. 3D), Bray-Curtis diversity for CORM2 samples was greater than SHI except for subject C (p<0.05), a common trend throughout this dataset. Therefore, these data suggest that saliva rather than blood-based medium potentially aids in plaque community formation more similar to *in vivo* SUPP.

### Model environmental factors minimally influence population diversity

While our model had broad similarity to initial donor microbiome composition, elevated *Neisseria* abundances led us to test if changing our incubation conditions might diminish its over-representation. We tested several conditions at 5 days in the model based on limiting *Neisseria* growth due to its aerobic and capnophilic nature. These included static incubation (T5nr), doubling the media replenishment rate (T5f2) and incubation without 5% CO2 (T5nc). For each condition, only the indicated change was made, and all others (CO2 status, rocking, temperature) unaltered. Unfortunately, we observed that these changes largely led to overgrowth of another taxa, *Lautropia mirabilis* (Fig. 4A, teal green). To assess if underlying composition of the SUPP microbiota were altered, we filtered both *Neisseria* and *L. mirabilis* reads and performed composition and diversity analyses (Fig. 4). Unfiltered read analyses are also available (see Supplemental 12A-E).

**Figure 4.**
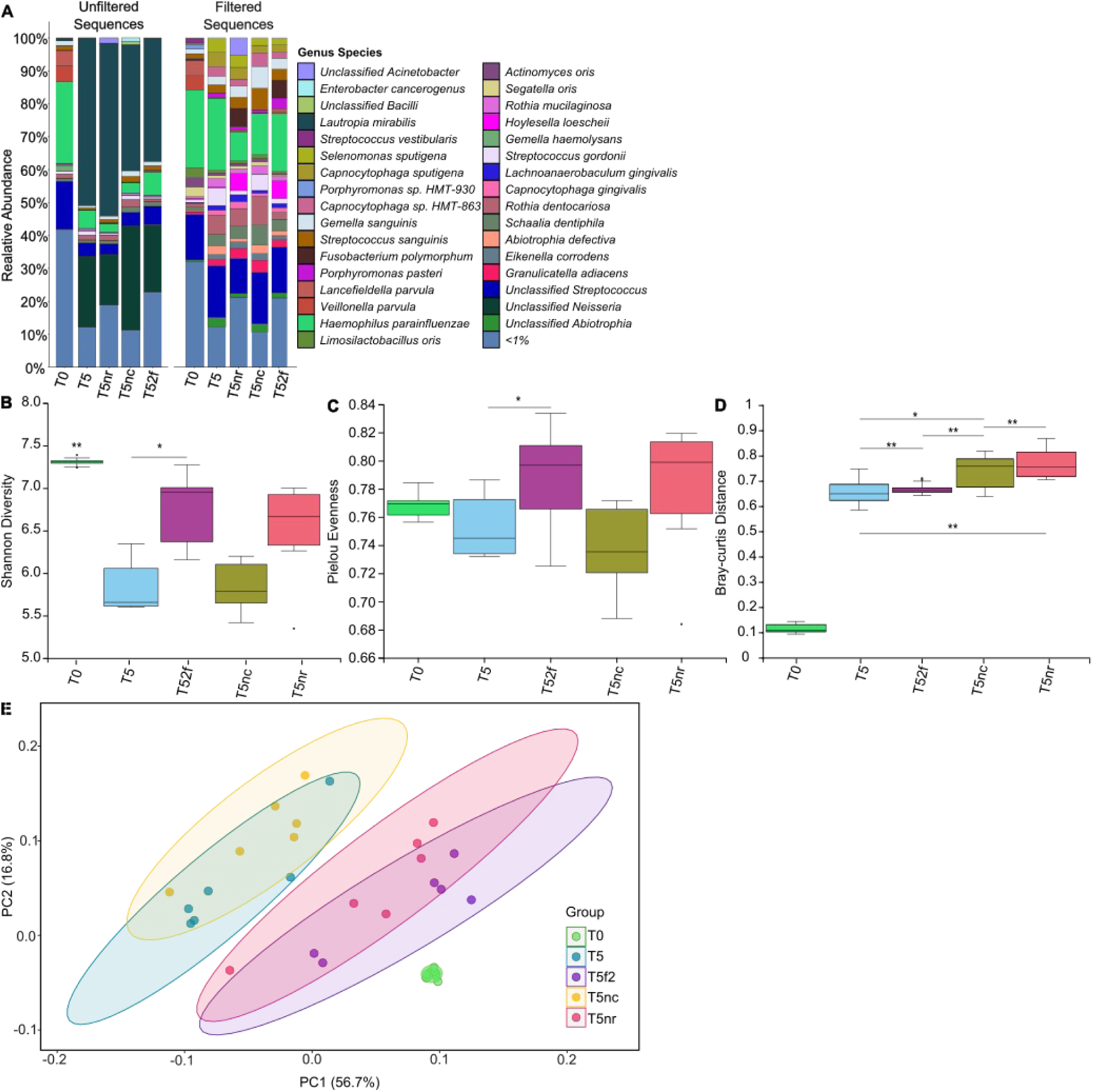
Physical parameters for model incubation affect microbiota composition. (A) DADA2 species assigned 16S profiles of the original donor SUPP (T0), initial day 5 testing conditions (T5), T5 incubated without rocking (T5nr), T5 incubated without 5% CO2 (T5nc), and T5 with 2x medium changes per day (T52f) revealed modest differences between conditions. Highly abundant *Neisseria* and *L. mirabilis* reads were computationally removed (Filtered Sequences) to reveal the underlying microbiota. Additional results for filtered and unfiltered comparisons are in Figs. S5-8, 12. (B) Shannon diversity and Pielou’s Evenness (C) indicate modest differences between conditions. β-diversity metrics for Bray-Curtis (E) and unweighted Unifrac comparisons (F) show similar results with T5f2 showing overall closer similarity to T0. α and β-diversity metrics utilized Kruskal-Wallis one-way ANOVA and pairwise PERMANOVA analyses respectively. n=6 for all comparisons, * p ≤ 0.05, ** p ≤ 0.01, and, *** p ≤ 0.001. Ellipses indicate a 99% confidence interval.

We preformed an analysis of composition of microbiomes with bias correction (ANCOM-BC) to determine if any of the incubation conditions affected the population of specific taxa (Supplemental S5-7). For static / non-rocking incubation (T5nr) we observed an expected increase of anaerobes including *Solobacterium moorei*, *Parvimonas* species and *Catonella morbi*. *C. matruchotii* was notably diminished in the T5 comparison (0.09% total abundance compared to the T0 at 0.26% total abundance). Doubling medium replenishment (T5f2) resulted in an increase of several facultative anaerobes including *Solobacterium moorei, Catonella morbi,* and *Fusobacterium polymorphum.* Interestingly, without CO_2_, we observed elevated amounts of *C. matruchotii* (0.25% to 0.31% relative abundance), *Schaalia* species*, Rothia dentocariosa* and *Streptococcus sanguinis*.

α-diversity analyses (Fig. 4B-E) reveal that diversity in more frequent medium exchange (T5f2) was greater than our original incubation conditions (T5), (p< 0.05); while static samples (T5nr) were not significantly different from twice fed samples. β-diversity metrics (Fig. 4D and 4E) interestingly showed that the greatest diversity compared to T0 plaque inoculum was our twice fed daily samples (T5f2), our static samples (T5nr), and our normal incubation conditions (T5). Based on these data, including our ANCOM-BC results, we believe that the most optimal *in vitro* incubation conditions to retain initial donor diversity was T5f2, and for future experiments to restrict duration to day 3 to limit overgrowth of SUPP taxa such as *Neisseria* and *L. mirabilis*.

### SUPP inoculum provides more similar in vitro populations vs saliva inoculum

Some plaque biofilm studies utilize unfiltered saliva as the source of inoculum (Edlund et al. 2013; Tian et al. 2010). While saliva contains SUPP species, it also contains species typically associated with other oral niches, particularly the tongue (Tamahara et al. 2025), and it is unknown how SUPP vs saliva inocula would affect *in vitro* community composition. Using our refined model conditions, we were able to assess the impact of inoculum source on SUPP composition (Fig. 5).

**Figure 5.**
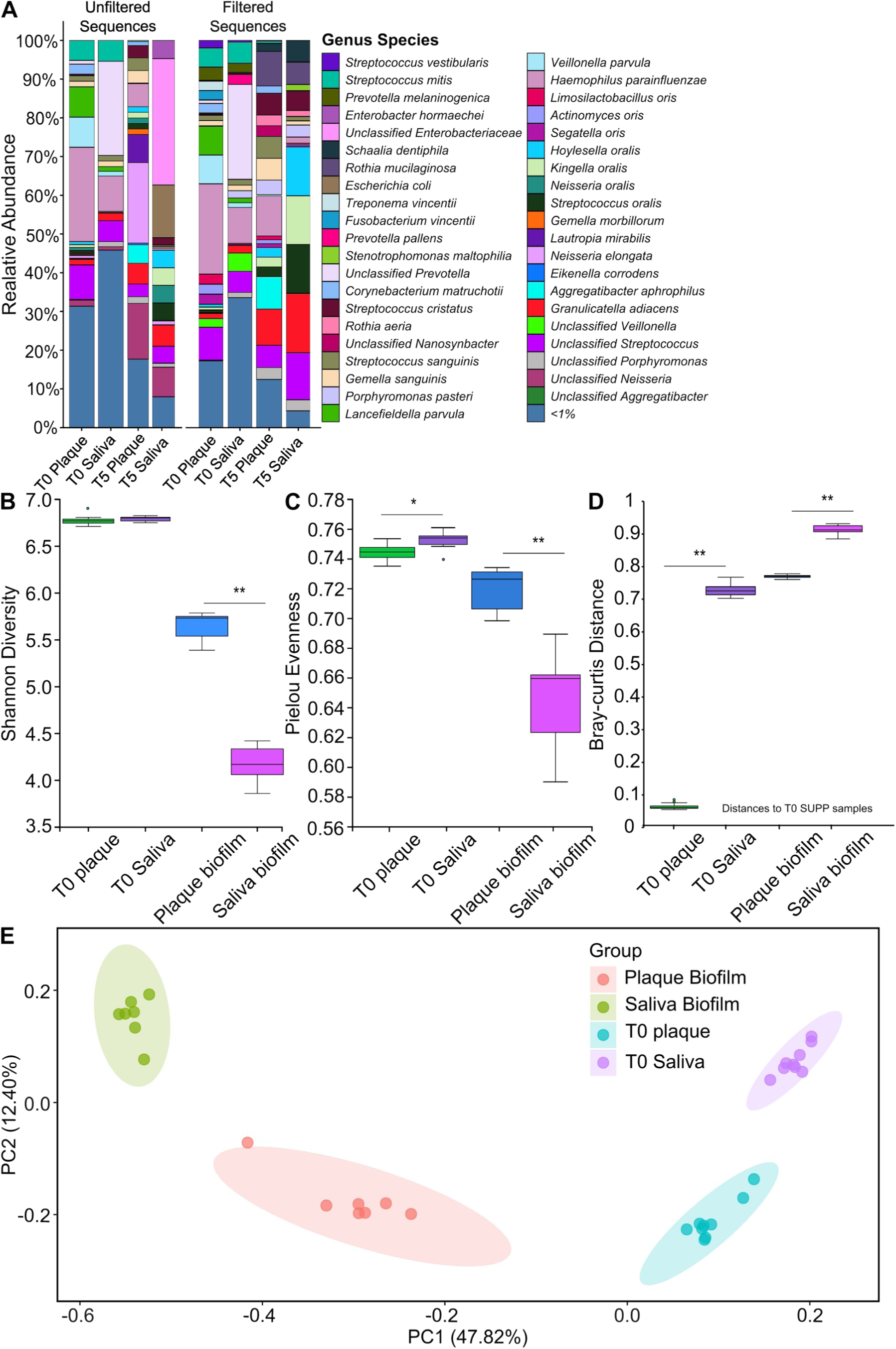
Inoculum source impacts model community composition. (A) DADA2 species assigned 16S profiles of donor SUPP (T0 Plaque) or donor unfiltered saliva (T0 Saliva) inoculums compared to 5 days after growth in the model (T5 Plaque/Saliva). Highly abundant *Neisseria*, *Enterobacteriaceae*, and *L. mirabilis* reads were computationally removed (Filtered Sequences) to reveal the underlying microbiota. Additional results for filtered and unfiltered comparisons are in Figs. S9, 13. (B) Shannon diversity and Pielou’s Evenness (C) indicate high similarity between SUPP and saliva inoculums and broad differences at T5 in the model for each. β-diversity metrics for Bray-Curtis (D) and unweighted Unifrac comparisons (E) reveal broad differences between each category with T5 model cultures exhibiting greater similarity to each other irrespective of inoculum source. α and β-diversity metrics utilized Kruskal-Wallis one-way ANOVA and pairwise PERMANOVA analyses respectively. n=7 for all comparisons, * p ≤ 0.05, ** p ≤ 0.01, and *** p ≤ 0.001. Ellipses indicate a 99% confidence interval.

As expected, initial community abundance profiles differed between SUPP and saliva inoculum (Fig. 5A). In saliva compared to SUPP we observed elevated amounts of *Prevotella* species (25% in saliva) and *Haemophilus parainfluenzae* (26% in SUPP vs 9% in saliva). We found increased abundance of the SUPP specialist *C. matruchotii* (∼2%, vs <1% in saliva) and elevated amounts of *Veillonella parvula* (∼8%, vs <1% in saliva). These differences support an expected founder effect which was reflected at later timepoints in the model. By day 3, *Haemophilus* is observed at 8% in SUPP inoculated samples compared to ∼1.5% in saliva-inoculated cultures. Likewise, *C. matruchotii* was no longer observed in saliva inoculated samples vs SUPP by day 3. Other species found primarily in saliva inoculum such as *Prevotella histicola* and *Prevotella pallens* were absent after 3 days, suggesting our model selects for specific organisms which would assimilate into communities more likely to be representative of dental plaque.

A weighted Unifrac analysis (Fig. 5E) reveals SUPP biofilms samples had clustered closer to the T0 plaque inoculate yet remain distinctly separate compared to the other communities based on their specific condition. This indicates these communities are different from one another as expected, where if saliva and plaque samples were the same, they would cluster much closer to one another as seen in the boiled vs filtered saliva PcoA data (Fig. 2F). Bray-Curtis dissimilarity metrics of day 3 samples for each inocula showed similar changes in diversity (Fig. 5C) yet SUPP-inoculated biofilms were more diverse than saliva (p<0.01 for all). All α-diversity metrics (Figure 5B, 5C and supplemental 9D-F) suggest SUPP-inoculated samples contained greater diversity in our model than saliva inoculated samples (p<0.05). Together, these data suggest using SUPP over saliva as an inoculum when focusing on SUPP related communities and interestingly show selection for SUPP-like microbiome signatures in the model irrespective of inoculum source.

### Validating mechanistic relationships within the SUPP model

With our model parameters decided we next wanted to test if we could quantify fitness for previously identified *in vitro* multispecies interactions between SUPP species. We specifically chose a phenotype that was moderate, to assess if our model had the ability to quantify even modest fitness changes. Previously, we described the ability of *C. matruchotii* to cross-feed on streptococcal-produced lactate (Almeida et al. 2023). We demonstrated lactate transfer between individual *C. matruchotii* and *Streptococcus mitis* cells (Puri et al. 2023) and showed that the *C. matruchotii lutABC* operon was largely responsible for lactate oxidation by this species. However, lactate oxidation was still evident in *C. matruchotii Δlut* mutants, likely due to multiple lactate dehydrogenases encoded on its genome. We were only able to show modest (∼2-fold) *C. matruchotii* fitness defects in *C. matruchotii Δlut -S. mitis* cocultures in defined medium under glucose limitation, which forced competition for glucose (i.e. mandated lactate cross-feeding for fitness) between these species.

To quantify fitness, we modified *C. matruchotii* wild type and *Δlut* strains to constitutively express *mCherry,* allowing us to measure fluorescence of these strains incorporated into *in vitro* SUPP biofilms via fluorescent microscopy over 3 days. We separately added fluorescent-expressing WT and *Δlut C. matruchotii* in a 1:1 ratio (by OD600) with SUPP inoculum and cultured each in parallel. Fluorescent *C. matruchotii* were readily visible in *in vitro* biofilms as were apparent “corncob” structures (Fig. 6A-B) reminiscent of those formed *in vivo* (Jessica Mark Welch 2022; Eren et al. 2014). After 3 days, we imaged cells mechanically debrided from separate biofilm discs (n=7 disc cultures per condition, ≥12 micrographs per culture) via fluorescence microscopy. Field of view acquisition was randomized and included only fields that had > 25% coverage with SUPP bacteria observed via light microscopy prior to fluorescence imaging. No visible fluorescence was evident in any SUPP control biofilm lacking labeled *C. matruchotii* using our acquisition settings. Figure 6C summarizes ImageJ (Schindelin et al. 2012) based total mCherry fluorescence quantification for all biofilm micrographs for each condition. We observed a small but significant (p<0.001) decrease (-1.7-fold) in abundance of the *C. matruchotii Δlut* mutant compared to the wild-type indicating its diminished fitness in a more complex medium and surrounding community than previously tested (Almeida et al. 2023). This demonstrates that the SUPP model can be used to successfully quantify the impact of even modest mechanistic phenotypes.

**Figure 6.**
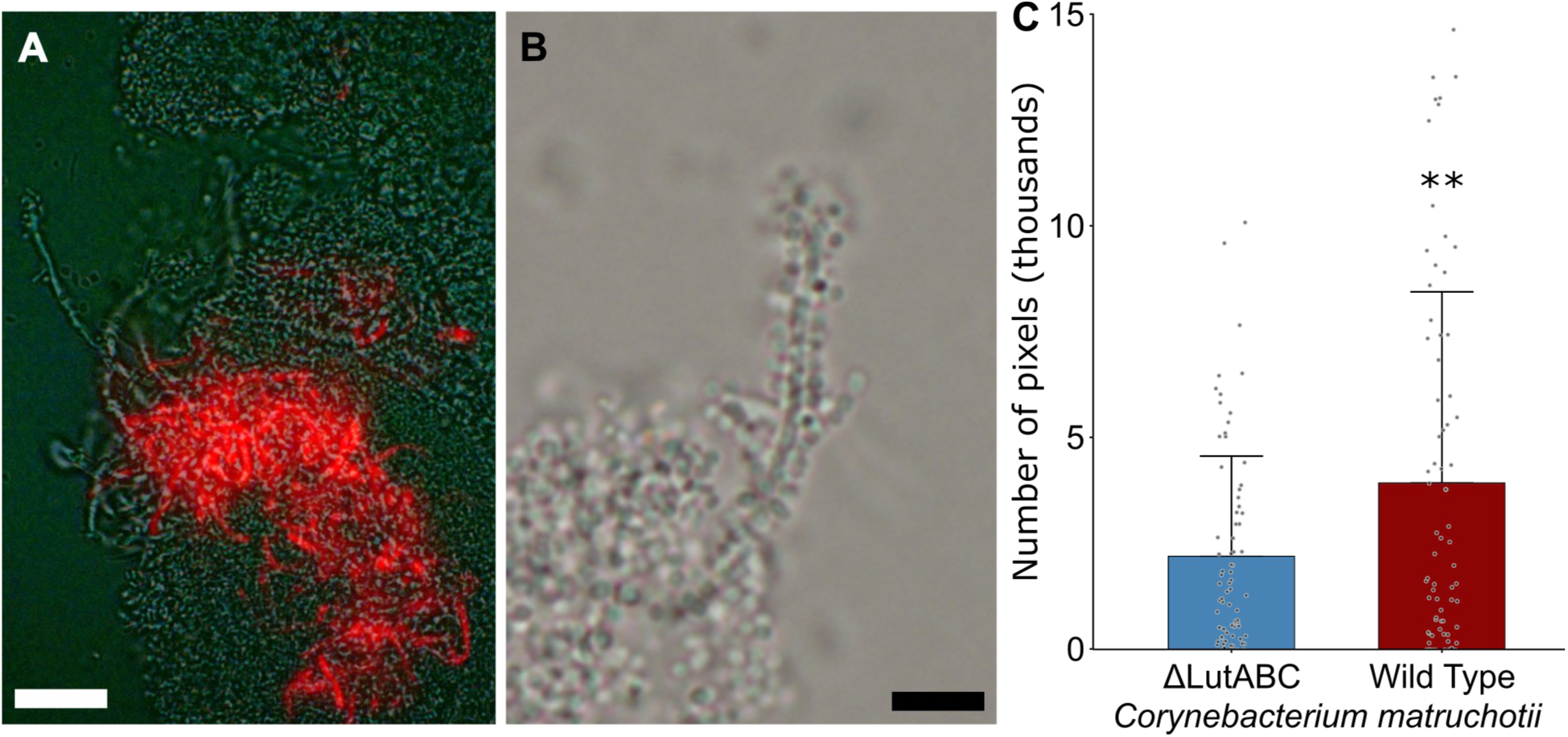
Impaired lactate metabolism in *C. matruchotii* reveals a fitness defect in SUPP model coculture. (A) mCherry-expressing *C. matruchotii* was easily visible via 400x epifluorescent microscopy in SUPP model biofilms 3-days post inoculum with donor SUPP. (B) Filamentous “corncob” structures were frequently observed via 1000x DIC microscopy. (C) Quantification of total mCherry observed in biofilm-containing fields of view (see methods) revealed a modest (∼1.7-fold) but significant fitness defect for a *C. matruchotii* mutant lacking the *lutABC* operon. n=7 indepdendent SUPP+*C. matruchotii* cultures for the WT and mutant strain each. 12 images were taken for each sample, with 84 images each for WT and mutant strains. Mcherry fluorescence was quantified via ImageJ (see methods). ** p ≤ 0.01 via Student’s t-test. Scalebars indicate 9 and 18 μm respectively.

## Discussion

We developed an *in vitro* model for the study of human supragingival plaque (SUPP) that mimics the host environment by focusing on utilization of a SUPP (vs saliva or handpicked consortia) inoculum, growth on saliva as a primary carbon and energy source, aerobic incubation, and hydroxyapatite (HA) as a biofilm substrate. We iteratively modified our model during development based on microbiome profiles at multi-day intervals compared to initial SUPP composition. This model is relatively low-cost and allows for large sample numbers, especially compared to *in vivo* models. It is also reductionist, allowing investigators to measure the impact of microbe-microbe and saliva composition changes independently of other host responses. We chose to develop this model due to prior *in vitro* models utilization of conditions that did not allow for robust growth of highly abundant SUPP commensal species. Many published models are quite adept at culturing oral pathogens and subgingival species at high abundance due to their anaerobic nature (Shapiro et al. 2002; Edlund et al. 2013; B. Guggenheim et al. 2001; Tsutsumi et al. 2018; Li et al. 2024), and / or utilization of specific consortia (B. Guggenheim et al. 2001; Shapiro et al. 2002), and rich media that do not utilize saliva as a primary source of carbon and energy (Edlund et al. 2013; Lamont et al. 2021). However, these models do not well reflect the native healthy SUPP population, necessitating this development.

There are numerous challenges to overcome in model development, and no *in vitro* model is a perfect mimic to *in vivo* growth. One challenge in model development is the growth medium. We tested several media formulations choosing our final medium (CORM2) based on its ability to cultivate SUPP bacteria while incorporating small amounts of other carbon and energy sources available *in vivo* (1mM sucrose, 0.5% human serum) to approximate dietary sugar and crevicular fluid exposure, in addition to 25% sterile saliva (Fig. 1). Using this, we were able to show the novel result that saliva sterilization methods (boiled vs filtration) had an unexpectedly minor impact on microbiome composition (Fig. 2). Another concern in model utilization, especially for microbiome assessment, is sample size. Many previous models were developed prior to current understandings on statistical power necessary to ascertain significant differences in microbiome composition (Utomo et al. 2024; P. Li et al. 2024; B. Guggenheim et al. 2001; Edlund et al. 2013; Blanc et al. 2014; Shapiro et al. 2002). This is further compounded by a large degree of inter-individual variation between donors (Huttenhower et al. 2012; Nearing et al. 2020) which we also observed in initial pilot studies with early model iterations (not shown). Because of this, we tested model parameters with a single SUPP donor before expanding further testing into multiple donors (Fig. 3) and utilized sample numbers from each individual sufficient to ensure a well powered study for most comparison metrics (Supplemental Discussion – power section). Volume of SUPP collection varied between donors but reliably allowed for ∼40 samples per donor.

With model conditions in place, we were able to assess how further environmental changes impacted growth of the initial SUPP population (Fig. 4). These differences can be utilized by researchers to further refine the model for cultivation of specific species of interest if desired. We tested these conditions in an attempt to limit observed overgrowth of some taxa, particularly *Neisseria* and *Lautropia* species. While altering conditions did little to abate this overgrowth, we suggest here to utilize intervals of 3 days to minimize it. Despite overgrowth, we were able to show that the remaining microbiota were highly consistent with those observed in donor SUPP (Figs. 3A, Supplemental X). Interestingly, *Neisseria* overgrowth (∼5-25% above initial abundance) was also observed in *in vivo* models using human dental materials in the human mouth (Tu et al. 2020; Utomo et al. 2024). Thus, this limitation is not unique to our model, where we are able to assess SUPP composition in a more cost-effective manner allowing for much higher sample numbers than can be tested in a human *in vivo* dental appliance. Another limitation of our model is that the presence of *Pseudomonas aeruginosa* in donor SUPP observed in ∼20% of donors (not shown) rapidly overtakes each culture, eliminating most remaining SUPP species. We recommend initial screening of donors by overnight aerobic culture of a toothpick SUPP scraping in LB broth. *P. aeruginosa-*containing donor cultures are readily apparent by pyocyanin (green) pigment production and these donors were excluded from the study.

We utilized the model to differentiate SUPP growth under different saliva preparation methods and compared *in vitro* microbiomes between differing inoculum sources (saliva vs SUPP) (Fig. 5). We were also able to add genetically manipulated commensal species to SUPP (Fig. 6) quantifying relatively small changes in fitness. These data verify that this model is sufficient to test commensal microbiota mechanisms including species-species metabolic interaction and biofilm formation properties between different strains. Further, our model will allow investigators to examine how saliva from different sources (eg, healthy vs diseased subjects) could influence the development of healthy SUPP. In future experiments we wish to further refine the model, and use it to compare metatranscriptome profiles of SUPP species from *in vivo* donors vs matched *in vitro* model populations to identify how closely microbial responses match gene expression in their native habitat. We will also utilize FISH-based microscopy methods to quantify established biogeographical relationships that we and others have identified (Mark Welch et al. 2016) in SUPP. This model will researchers to expand initial *in vitro* coculture interactions into a more relevant and complex polymicrobial context without costly *in vivo* experimental intervention revealing the impact of bacterial interactions separate from host influences, further refining the overall study of the impact of the commensal microbiota on host health.

## Materials and Methods

### Saliva collection

Saliva was collected according to the University of Louisville IRB protocol #24.0794, reference #794353. 40-50 mL of stimulated saliva was collected from donors who did not eat 2 hours prior to collection, had not undergone antibiotic exposure for >6 months, and who had no self-reported current illnesses. Saliva flow was stimulated by mastication on pieces of Parafilm® M (Bemis Company, Inc.) and frozen at -20 °C until processed.

### Saliva preparation and sterilization

Saliva was thawed overnight at 4°C. <u>Filter sterilization</u> was performed using methods from (De Jong and Van Der Hoeven 1987). Saliva was pooled from a minimum of 3 donors and DL-Dithiothreitol (DDT) (Thermo Scientific) was added to 2.5mM and mixed at 4°C for 10 m. Saliva was then centrifuged at 4,000xg for 1hr at 4 °C separating saliva from an insoluble pellet of cell debris. Supernatants were then either filter sterilized through a 0.22μm PES 50mL Centrifuge Tube Filter System (CELLTREAT 229710) or heat sterilized via immersion into a boiling water bath for 15 m and aliquots frozen at -20 °C.

### Media formulation

SHI medium: proteose peptone 10 g/L; trypticase peptone 5.0 g/L; yeast extract 5.0 g/L; KCl 2.5 g/L; sucrose 5 g/L; haemin 5 mg/L; VitK 1 mg/L; urea 0.06 g /L, arginine 0.174 g/L; mucin (type III, porcine, gastric) 2.5 g/L; sheep blood 5% and N -acetylmuramic acid (NAM) 10 mg/L.

<u>CORM</u>: (Teknova EZ_RICH (Teknova, USA) with saliva and other amendments): 50 mL -EZ rich defined media (Mops buffer 40 mM, Potassium phosphate dibasic 1.32 mM, 10X ACGU solution 5mL, 5X Supplement EZ solution 10mL), Lipoic acid 484 nM, Folic acid 2.26 uM, NAD 4.21 nM, Riboflavin 2.65 nM, 1000x amino acid + nucleotide block (see table below) 50 μL, Vitamin block (see table in Supplemental) 50 μL. Type I water 28.25 mL.

<u>CORM2</u>: 40 mL – 100% pooled saliva, 10mL. 10X MOPS buffer (400mM MOPS) (Teknova, Inc), 4 mL. 1% Yeast extract (w/v), 4 mL. 100% heat inactivated human serum, 200 μL. 100 mM sucrose, 100 μL. <u>All media were if needed, adjusted to pH 7.5</u> prior to filter sterilization.

### Hydroxyapatite (HA) disc treatment for saliva pellicle formation

HA discs (12.7mm x 3.81 mm, Biosurface Technologies Co, Bozeman, MT USA) were placed in a glass Petri dish and autoclave sterilized. Filtered saliva was diluted to 25% in 1x PBS (v/v%), sealed with Parafilm® and placed in a 37°C 5% CO2 incubator for 24 h before use. Sterile tweezers were used to add saliva-coated HA discs to each well in a sterile polystyrene 24-well microassay plate.

### Ex vivo SUPP collection

Supragingival plaque was collected according to protocol# 24.0794, approved by the University of Louisville IRB. Donors who did not take antibiotics within >6 months and had no self-reported illness were used. Donors were asked to refrain from oral hygiene practices for 24 hours prior to collection, and were asked to refrain from drinking, smoking, eating or vaping 2 hours prior to collection. Upon arrival donors were given sterile wood toothpicks and self-collected SUPP after instruction shown to avoid sampling subgingival plaque by placing the toothpick above the gumline and crevices of teeth and pulling away from the gums. After each scrape of the tooth/crevices between teeth, biofilms on each toothpick were placed in a 1.5mL microcentrifuge tube with 1mL PBS until tubes were filled with ∼0.1 mL loose precipitated SUPP, donors used 1 toothpick per scrape. All teeth were used to collect SUPP. Plaque was immediately used after collection and inoculated into medium at an OD of 0.05 (OD600nm), remaining SUPP was centrifuged at 12,000 x g for 5 min, supernatant discarded, and the SUPP pellet frozen at -20 °C for evaluation of T0 SUPP samples.

### SUPP model assembly

Saliva coated hydroxyapatite (sHA) discs were placed in 24 well plates and filled with 350 µl of medium each. Wells containing sHA discs were seeded with plaque diluted into the medium at an OD600 of 0.05 alongside at least 1 sHA disc in sterile medium as a contamination control. Each plate was placed in a 37 °C incubator at 5% CO2 on a rocking platform incubator unless indicated otherwise (see section on environmental changes). Once daily, (or twice for environmental change testing) each sHA disc was aseptically washed in 3 separate wells containing 800 µl 1x PBS to remove loosely attached / planktonic cells. After washing, sHA discs were placed into new wells containing sterile pre-warmed media.

### In vitro SUPP collection

At indicated timepoints, sHA discs were washed and discs were placed into sterile wells containing 400µl of 1x PBS. The washed disc was placed under a dissecting microscope, and sterile toothpicks were used to remove attached biofilm material. SUPP-containing buffer solution was transferred to a 1.5mL microcentrifuge tube and spun at 12,000 x g for 5 m. Supernatant was discarded and each pellet was frozen at -20 °C until DNA isolation.

### DNA isolation from SUPP

DNA was extracted using the MasterPure™ Complete DNA & RNA Purification Kit (Lucigen / LGC Biosearch Technologies). Cell lysis was accomplished by addition of cell pellets in 300 μL of lysis solution containing Proteinase K into Lysing Matrix B tubes (MP Biomedicals-1169110-CF) and incubation at 55 °C for 15m prior to beadbeating. Cells were incubated in a Omni Bead Ruptor 24 for 1 min at 6 m/s, and incubated on ice for 3m and repeated once more. DNA precipitation and cleaning were done following the manufacturer’s instructions and DNA pellets were resolubilized in 40 μL of TE buffer. DNA concentration and purity were measured via UV spectroscopy in a BioTek Take3 micro-volume plate (Agilent BioTek).

### V1-V2 16S PCR amplification

Universal 16s rRNA primer V1 (27f-YM – AGAGTTTGATYMTGGCTCAG) and V2 ( 338R-R TGCTGCCTCCCGTAGRAGT) (W.G. Wade and Prosdocimi 2020) were used to amplify 16s rRNA (∼311 bp amplicon size) using Q5 HiFi mastermix (New England Biolabs, USA) in a 25 µl reaction (98 °C 1 m, 98 °C 10 s, 55 °C 30 s, 72 °C 30s, for 18 cycles, 72 °C 1 min, 4 °C hold). Amplicon size and integrity was confirmed by electrophoresis on 2% agarose gels stained with SYBR-Safe (Thermo-Fisher, USA). Sequencing was performed at the University of Rhode Island Genomics and Sequencing Center (Kingston, RI) or the University of Louisville (Louisville, KY) sequencing technology center on an Illumina MiSeq using 2×250bp paired end sequencing.

### Microbiome composition analysis

Sequences were imported using QIIME2-2025.4 (Bolyen et al. 2019). Denoising and determination of amplicon sequence variants (ASVs) were done using the DADA2 plugin to join paired-end sequences, performing quality filtering, and chimera checking. Each data set was trimmed accordingly based on the quality of the sequences. α and β-diversity analyses were done using QIIME2 based on the sample depth from each dataset’s rarefaction curve. In most cases native QIIME2 outputs were redrawn in RStudio. α-diversity of our samples was assessed via Shannon, Faith PD, Pielou eveness, and Chao1 metrics. β-diversity was assessed via weighted Unifrac and Bray-Curtis Dissimilarity matrices. Each α-diversity graph calculated significance from a Kruskal-Wallis test and Bray-Curtis utilized a PERMANOVA test with corrected-p values respectively. Analysis of Composition of Microbiomes with Bias Correction (ANCOM-BC) (Lin and Peddada 2020) was performed on saliva and boiled saliva samples, the threshold for significance was α=0.0001. All scripts used for sequencing and bioinformatics analyses are present in Supplemental. All sequences in this used in this study can be found in the NCBI SRA database under bioproject PRJNA1256350.

### Fluorescence labeling of *C. matruchotii* for fitness testing

We used the replicating pCGL243 vector (Almeida et al. 2023) as a backbone to add a constitutive *xylR* promoter upstream of *mCherry* to create the fluorescence expression plasmid pSO002. This vector was transformed into both *C. matruchotii* ATCC 14266 and *C. matruchotii ΔlutABC* generated previously (Almeida et al. 2023). Transformations were carried out via electroporation as previously described (Almeida et al. 2023) and transformants were selected for on BHI-YE + 10 μg/mL kanamycin plates and incubated for 4 d at 37°C until transformants were visible. mCherry production was confirmed via epifluorescence microscopy and all strains showed roughly uniform per-cell fluorescence in red but not green or blue channel filters.

### Incorporation of *C. matruchotii* into the SUPP model

All *C. matruchotii* strains were cultured at 37 °C on BHI-YE + 10 μg/mL kanamycin broth for 3 days under standard incubation conditions. Following incubation, *C. matruchotii* cultures were washed three times with 1× PBS to remove residual medium. Optical densities (ODs) were then measured, and washed *C. matruchotii* cells were adjusted to a 1:1 ratio with plaque samples (by OD600) for subsequent experiments.

### Imaging and analysis

Biofilm cells for imaging were prepared by toothpick debridement from each sHA disc, debrided SUPP quantified by spectrophotometry, and the total SUPP concentration normalized to an identical optical density for each sample prior to imaging. We imaged mCherry fluorescent *C. matruchotii* using a Nikon Eclipse E600 epifluorescence microscope at 40x/0.75 magnification through a 590-650 nm TX RED filter using a Nikon C-SHG1 Super High Pressure Mercury Lamp. The images were taken by a Nikon DS-Ri1 camera and recorded by Nikon NIS-Elements F software. Exposure was set to 4ms with an analog gain of 2x. To obtain random images without bias, the initial field of view was imaged and then moved roughly 2 cardinal directions without prior viewing of the specific field under fluorescence for each photo taken. 168 photos were taken between the two conditions, 12 photos per sample, 84 photos per condition. Before each image was taken we ensured via phase contrast microscopy that at least 25% of each field of view contained bacterial cells ensure that each image had biofilm cells with or without fluorescence and that our quantification was not biased by oversampling of empty fields of view. If <25% of the field of view contained cells we moved to the adjacent field of view repeating the process above until 12 photos per sample were taken.

Photos were analyzed using ImageJ to quantify all red pixels per image. Images were captured only when the focused visible light field of view contained ≥20% biofilm biomass, ensuring no empty fields of view were acquired. No red-filter fluorescence was observed in SUPP-only controls under our acquisition settings. Photos were quantified only for the red channel; background was subtracted via the ImageJ “subtract background” function choosing the rolling paraboloid option with a radius of 75. Thresholds were set from 50 to 225. The photos were processed via the smooth option and despeckled. Images were then converted to a binary mask. Number of pixels within the photos were then quantified Outlier values were removed from each group based on each groups lower and upper fence. Outliers were calculated using the 1.5IQR rule, where Q1-1.5 (interquartile range) was used to find the lower fence and Q3+1.5 (interquartile range) was used to find the upper fence. Total pixel values >10744 were determined to be outliers. 5 samples in the WT condition were excluding for reaching above the upper fence, and 8 samples were removed from the *ΔlutABC* samples for reaching above the upper fence. A student’s T-test was applied to determine significance between conditions.

## Supporting information

Supplemental figures, methods and references

## Work Cited

Adami, Guy R., Wei Li, Stefan J. Green, Elissa M. Kim, and Christine D. Wu. 2025. “Ex Vivo Oral Biofilm Model for Rapid Screening of Antimicrobial Agents Including Natural Cranberry Polyphenols.” Scientific Reports 15 (1): 6130. 10.1038/s41598-025-87382-0.

Almeida, Eric, Surendra Puri, Alex Labossiere, Subashini Elangovan, Jiyeon Kim, and Matthew Ramsey. 2023. “Bacterial Multispecies Interaction Mechanisms Dictate Biogeographic Arrangement between the Oral Commensals Corynebacterium Matruchotii and Streptococcus Mitis.” mSystems 8 (5): e00115–23. 10.1128/msystems.00115-23.

Bao, Kai, Adam Papadimitropoulos, Baki Akgül, Georgios N. Belibasakis, and Nagihan Bostanci. 2015. “Establishment of an Oral Infection Model Resembling the Periodontal Pocket in a Perfusion Bioreactor System.” Virulence 6 (3): 265–73. 10.4161/21505594.2014.978721.

Blanc, V., S. Isabal, M. C. Sánchez, et al. 2014. “Characterization and Application of a Flow System for *in Vitro* Multispecies Oral Biofilm Formation.” Journal of Periodontal Research 49 (3): 323–32. 10.1111/jre.12110.

Bolyen, Evan, Jai Ram Rideout, Matthew R. Dillon, et al. 2019. “Reproducible, Interactive, Scalable and Extensible Microbiome Data Science Using QIIME 2.” Nature Biotechnology 37 (8): 852–57. 10.1038/s41587-019-0209-9.

Bradshaw, D. J., P. D. Marsh, K. M. Schilling, and D. Cummins. 1996. “A Modified Chemostat System to Study the Ecology of Oral Biofilms.” Journal of Applied Bacteriology 80 (2): 124–30. 10.1111/j.1365-2672.1996.tb03199.x.

Caufield, P. W., C. N. Schön, P. Saraithong, Y. Li, and S. Argimón. 2015. “Oral Lactobacilli and Dental Caries.” Journal of Dental Research 94 (9 Suppl): 110S–118S. 10.1177/0022034515576052.

De Jong, M. H., and J. S. Van Der Hoeven. 1987. “The Growth of Oral Bacteria on Saliva.” Journal of Dental Research 66 (2): 498–505. 10.1177/00220345870660021901.

Deari, Shengjile, Monika Gothwal, Kay Gränicher, Thomas Thurnheer, Thomas Attin, and Lamprini Karygianni. 2025. “The Effect of Pellicle on Biofilm Formation in a Supragingival Biofilm Model.” Clinical and Experimental Dental Research 11 (6): e70276. 10.1002/cre2.70276.

Dini, Caroline, Raphael Cavalcante Costa, Martinna Bertolini, et al. 2023. “In-Vitro Polymicrobial Oral Biofilm Model Represents Clinical Microbial Profile and Disease Progression during Implant-Related Infections.” Journal of Applied Microbiology 134 (11): lxad265. 10.1093/jambio/lxad265.

Edlund, Anna, Youngik Yang, Adam P. Hall, et al. 2013. “An in Vitrobiofilm Model System Maintaining a Highly Reproducible Species and Metabolic Diversity Approaching That of the Human Oral Microbiome.” Microbiome 1 (1): 25. 10.1186/2049-2618-1-25.

Eren, A. Murat, Gary G. Borisy, Susan M. Huse, and Jessica L. Mark Welch. 2014a. “Oligotyping Analysis of the Human Oral Microbiome.” Proceedings of the National Academy of Sciences of the United States of America 111 (28): E2875– 84. 10.1073/pnas.1409644111.

Eren, A. Murat, Gary G. Borisy, Susan M. Huse, and Jessica L. Mark Welch. 2014b. “Oligotyping Analysis of the Human Oral Microbiome.” Proceedings of the National Academy of Sciences of the United States of America 111 (28): E2875–2884. 10.1073/pnas.1409644111.

Ghesquière, Justien, Kenneth Simoens, Erin Koos, Nico Boon, Wim Teughels, and Kristel Bernaerts. 2023. “Spatiotemporal Monitoring of a Periodontal Multispecies Biofilm Model: Demonstration of Prebiotic Treatment Responses.” Applied and Environmental Microbiology 89 (10): e01081–23. 10.1128/aem.01081-23.

Giertsen, E., R. A. Arthur, and B. Guggenheim. 2011. “Effects of Xylitol on Survival of Mutans Streptococci in Mixed-Six-Species in Vitro Biofilms Modelling Supragingival Plaque.” Caries Research 45 (1): 31–39. 10.1159/000322646.

Guggenheim, B., E. Giertsen, P. Schüpbach, and S. Shapiro. 2001. “Validation of an in Vitro Biofilm Model of Supragingival Plaque.” Journal of Dental Research 80 (1): 363–70. 10.1177/00220345010800011201.

Guggenheim, Bernhard, Merlin Guggenheim, Rudolf Gmür, Elin Giertsen, and Thomas Thurnheer. 2004. “Application of the Zürich Biofilm Model to Problems of Cariology.” Caries Research 38 (3): 212–22. 10.1159/000077757.

Guggenheim, M., S. Shapiro, R. Gmür, and B. Guggenheim. 2001. “Spatial Arrangements and Associative Behavior of Species in an In Vitro Oral Biofilm Model.” Applied and Environmental Microbiology 67 (3): 1343–50. 10.1128/AEM.67.3.1343-1350.2001.

Hajishengallis, G., and R. J. Lamont. 2012. “Beyond the Red Complex and into More Complexity: The Polymicrobial Synergy and Dysbiosis (PSD) Model of Periodontal Disease Etiology.” Molecular Oral Microbiology 27 (6): 409–19. 10.1111/j.2041-1014.2012.00663.x.

Hajishengallis, George, Richard P. Darveau, and Michael A. Curtis. 2012. “The Keystone Pathogen Hypothesis.” Nature Reviews. Microbiology 10 (10): 717–25. 10.1038/nrmicro2873.

Huttenhower, Curtis, Dirk Gevers, Rob Knight, et al. 2012. “Structure, Function and Diversity of the Healthy Human Microbiome.” Nature 486 (7402): 207–14. 10.1038/nature11234.

Jessica Mark Welch. 2022. “Biogeography of a Human Oral Microbiome at the Micron Scale | PNAS.” September 29. https://www.pnas.org/doi/10.1073/pnas.1522149113?cookieSet=1.

Keevil, C. W., D. J. Bradshaw, A. B. Dowsett, and T. W. Feary. 1987. “Microbial Film Formation: Dental Plaque Deposition on Acrylic Tiles Using Continuous Culture Techniques.” Journal of Applied Bacteriology 62 (2): 129–38. 10.1111/j.1365-2672.1987.tb02390.x.

Kwon, TaeHyun, Ira B. Lamster, and Liran Levin. 2021. “Current Concepts in the Management of Periodontitis.” International Dental Journal 71 (6): 462–76. 10.1111/idj.12630.

Lamont, Eleanor I., Archita Gadkari, Kristopher A. Kerns, et al. 2021. “Modified SHI Medium Supports Growth of a Disease-state Subgingival Polymicrobial Community in Vitro.” Molecular Oral Microbiology 36 (1): 37–49. 10.1111/omi.12323.

Li, Pengpeng, Yuwen Zhang, Dongru Chen, and Huancai Lin. 2024. “Investigation of a Novel Biofilm Model Close to the Original Oral Microbiome.” Applied Microbiology and Biotechnology 108 (1): 330. 10.1007/s00253-024-13149-8.

Lin, Huang, and Shyamal Das Peddada. 2020. “Analysis of Compositions of Microbiomes with Bias Correction.” Nature Communications 11 (1): 3514. 10.1038/s41467-020-17041-7.

Loesche, W. J., D. R. Bradbury, and M. P. Woolfolk. 1977. “Reduction of Dental Decay in Rampant Caries Individuals Following Short-Term Kanamycin Treatment.” Journal of Dental Research 56 (3): 254–65. 10.1177/00220345770560031101.

Mark Welch, Jessica L., Blair J. Rossetti, Christopher W. Rieken, Floyd E. Dewhirst, and Gary G. Borisy. 2016. “Biogeography of a Human Oral Microbiome at the Micron Scale.” Proceedings of the National Academy of Sciences 113 (6): E791–800. 10.1073/pnas.1522149113.

Marsh, P. D. 1994. “Microbial Ecology of Dental Plaque and Its Significance in Health and Disease.” Advances in Dental Research 8 (2): 263–71. 10.1177/08959374940080022001.

Marsh, Philip D. 2006. “Dental Plaque as a Biofilm and a Microbial Community – Implications for Health and Disease.” BMC Oral Health 6 (Suppl 1): S14. 10.1186/1472-6831-6-S1-S14.

Naginyte, Monika, Thuy Do, Josephine Meade, Deirdre Ann Devine, and Philip David Marsh. 2019. “Enrichment of Periodontal Pathogens from the Biofilms of Healthy Adults.” Scientific Reports 9 (1): 5491. 10.1038/s41598-019-41882-y.

Nearing, Jacob T., Vanessa DeClercq, Johan Van Limbergen, and Morgan G. I. Langille. 2020. “Assessing the Variation within the Oral Microbiome of Healthy Adults.” mSphere 5 (5): 10.1128/msphere.00451-20. 10.1128/msphere.00451-20.

Perera, Dasith, Anthony McLean, Viviana Morillo-López, et al. 2022. “Mechanisms Underlying Interactions between Two Abundant Oral Commensal Bacteria.” The ISME Journal 16 (4): 948–57. 10.1038/s41396-021-01141-3.

Pigman, Ward, Howard C. Elliott, and R. O. Laffre. 1952. “An Artificial Mouth for Caries Research.” Journal of Dental Research 31 (5): 627–33. 10.1177/00220345520310050501.

Puri, Surendra R., Eric Almeida, Subhashini Elangovan, et al. 2023. “Mechanistic Assessment of Metabolic Interaction between Single Oral Commensal Cells by Scanning Electrochemical Microscopy.” Analytical Chemistry 95 (22): 8711–19. 10.1021/acs.analchem.3c01498.

Ramsey, Matthew M., and Marvin Whiteley. 2009. “Polymicrobial Interactions Stimulate Resistance to Host Innate Immunity through Metabolite Perception.” Proceedings of the National Academy of Sciences of the United States of America 106 (5): 1578–83. 10.1073/pnas.0809533106.

Rathee, Manu, and Amit Sapra. 2025. “Dental Caries.” In *S*tatPearls. StatPearls Publishing. http://www.ncbi.nlm.nih.gov/books/NBK551699/.

Russell, C., and W. A. Coulter. n.d. Continuous Monitoring of pH and Eh in Bacterial Plaque Grown on a Tooth in an Artificial Mouth.

Schindelin, Johannes, Ignacio Arganda-Carreras, Erwin Frise, et al. 2012. “Fiji: An Open-Source Platform for Biological-Image Analysis.” Nature Methods 9 (7): 676 –82. 10.1038/nmeth.2019.

Shapiro, S., E. Giertsen, and B. Guggenheim. 2002. “An in Vitro Oral Biofilm Model for Comparing the Efficacy of Antimicrobial Mouthrinses.” Caries Research 36 (2): 93–100. 10.1159/000057866.

Spatafora, Grace, Yihong Li, Xuesong He, Annie Cowan, and Anne C. R. Tanner. 2024. “The Evolving Microbiome of Dental Caries.” Microorganisms 12 (1): 121. 10.3390/microorganisms12010121.

Tamahara, Toru, Atsumu Kouketsu, Satoshi Fukase, et al. 2025. “Ecological and Functional Landscape of the Oral Microbiome: A Multi-Site Analysis of Saliva, Dental Plaque and Tongue Coating.” Microorganisms 14 (1): 2. 10.3390/microorganisms14010002.

Tian, Y., X. He, M. Torralba, et al. 2010. “Using DGGE Profiling to Develop a Novel Culture Medium Suitable for Oral Microbial Communities.” Molecular Oral Microbiology 25 (5): 357–67. 10.1111/j.2041-1014.2010.00585.x.

Tsutsumi, K., M. Maruyama, A. Uchiyama, and K. Shibasaki. 2018. “Characterisation of a Sucrose-Independent in Vitro Biofilm Model of Supragingival Plaque.” Oral Diseases 24 (3): 465–75. 10.1111/odi.12779.

Tu, Yan, Yuan Wang, Lingkai Su, Beibei Shao, Zhuhui Duan, and Shuli Deng. 2020. “In Vivo Microbial Diversity Analysis on Different Surfaces of Dental Restorative Materials via 16S rDNA Sequencing.” Medical Science Monitor : International Medical Journal of Experimental and Clinical Research 26 (July): e923509-1–e923509-11. 10.12659/MSM.923509.

Utomo, Romualdus Nugraha Catur, Alena Lisa Palkowitz, Lin Gan, et al. 2024a. “In Vitro Plaque Formation Model to Unravel Biofilm Formation Dynamics on Implant Abutment Surfaces.” Journal of Oral Microbiology 16 (1): 2424227. 10.1080/20002297.2024.2424227.

Utomo, Romualdus Nugraha Catur, Alena Lisa Palkowitz, Lin Gan, et al. 2024b. “In Vitro Plaque Formation Model to Unravel Biofilm Formation Dynamics on Implant Abutment Surfaces.” Journal of Oral Microbiology 16 (1): 2424227. 10.1080/20002297.2024.2424227.

Wade, W.G., and E. M. Prosdocimi. 2020. “Profiling of Oral Bacterial Communities.” Journal of Dental Research 99 (6): 621–29. 10.1177/0022034520914594.

Wade, William G. 2021. “Resilience of the Oral Microbiome.” Periodontology 2000 86 (1): 113–22. 10.1111/prd.12365.

Wang, Cong, Tian Xu, Chaminda Jayampath Seneviratne, Louis Jun Ye Ong, and Yinghong Zhou. 2024. “Modelling Periodontitis in Vitro: Engineering Strategies and Biofilm Model Development.” Frontiers in Biomaterials Science 3 (July). 10.3389/fbiom.2024.1380153.

Zhang, Yixin, Jiakun Fang, Jingyi Yang, et al. 2022. “Streptococcus Mutans-Associated Bacteria in Dental Plaque of Severe Early Childhood Caries.” Journal of Oral Microbiology 14 (1): 2046309. 10.1080/20002297.2022.2046309.

