## Supplemental figures, methods and references for "“What’s SUPP” developing an *in vitro* model for healthy oral biofilms"

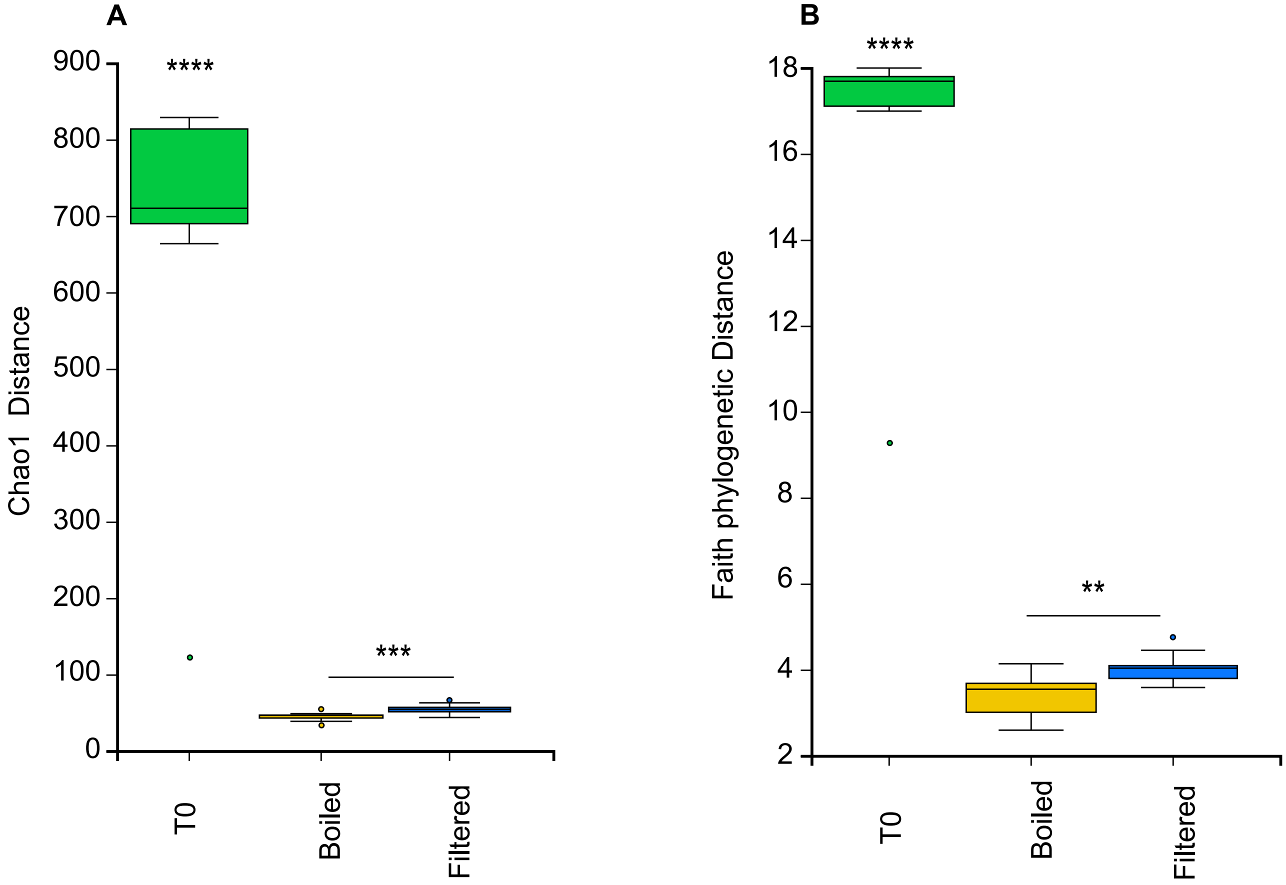


**Figure S1.** See Figure 2 for remaining dataset. (A) Chao1 Distance, measurement of total species richness based on the abundance of rare taxa within a system, (B) Faith phylogenetic distance alpha diversity measured by total sum of branch length of all branches within the tree that spans the members found within the sample. Calculations were done by qiime2 via Kruskal–Wallis test (one-way ANOVA) for alpha diversity where * denotes $p \leq 0.05$, ** denotes $p \leq 0.01$, *** denotes $p \leq 0.001$ and **** denotes $p \leq 0.0001$

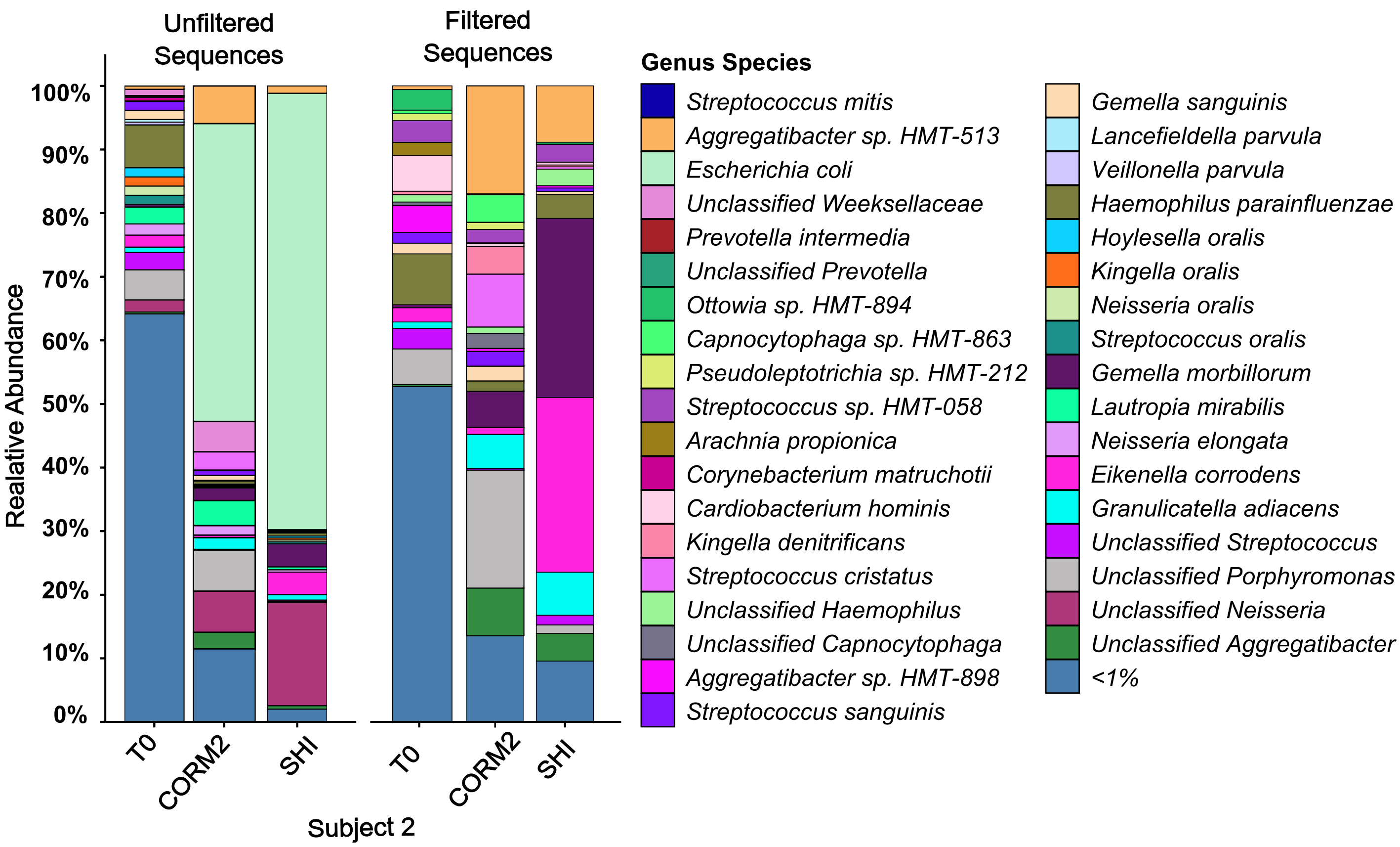


**Figure S2.** See figure 3 for associated data. Total percent abundance bar graph of identified species within samples (n=7 per bar graph) grown in CORM2 media compared to samples grown in SHI media post 3 days. “Unfiltered” refers to dataset where ASV’s of Genus such as Neisseria (magenta), E. coli (light green) and Lautropia mirabilis (dark purple) are present and compared to the same dataset where these listed ASV’s have been computationally “filtered out”. Bar graphs show underlying diversity is still present when these sequences are filtered out of the dataset.

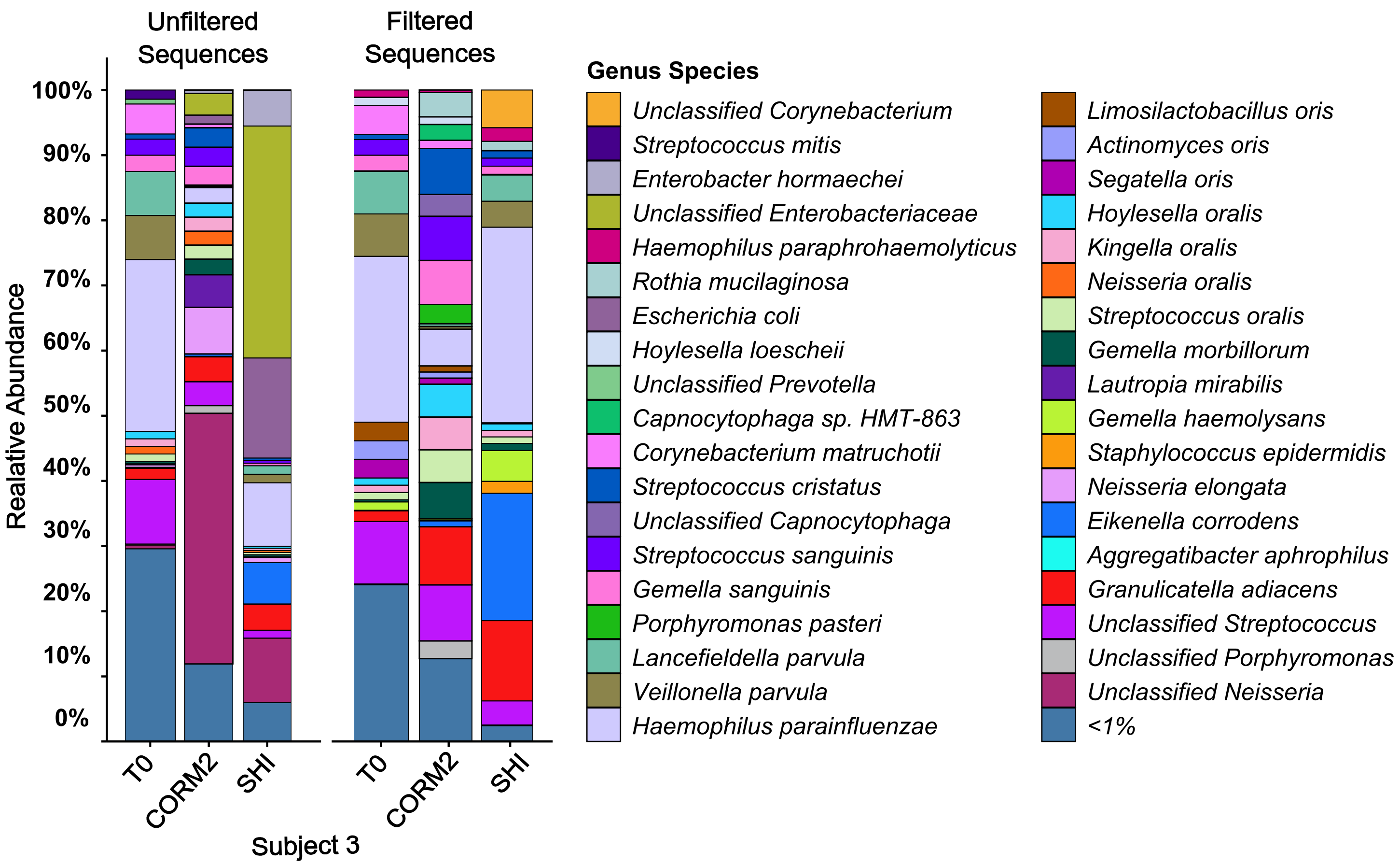


**Figure S2.** See figure 3 for associated data. Total percent abundance bar graph of identified species within samples (n=7 per bar graph) were grown in CORM2 media compared to samples grown in SHI media post 3 days. “Unfiltered” refers to dataset where ASV’s belonging to genus Neisseria (magenta), Enterobacteriaceae ( dark yellow and dark gray) and Lautropia mirabilis (dark purple) are present and compared to the same dataset where these listed ASV’s have been computationally “filtered out”. Bar graphs show underlying diversity is still present when these sequences are filtered out of the dataset.

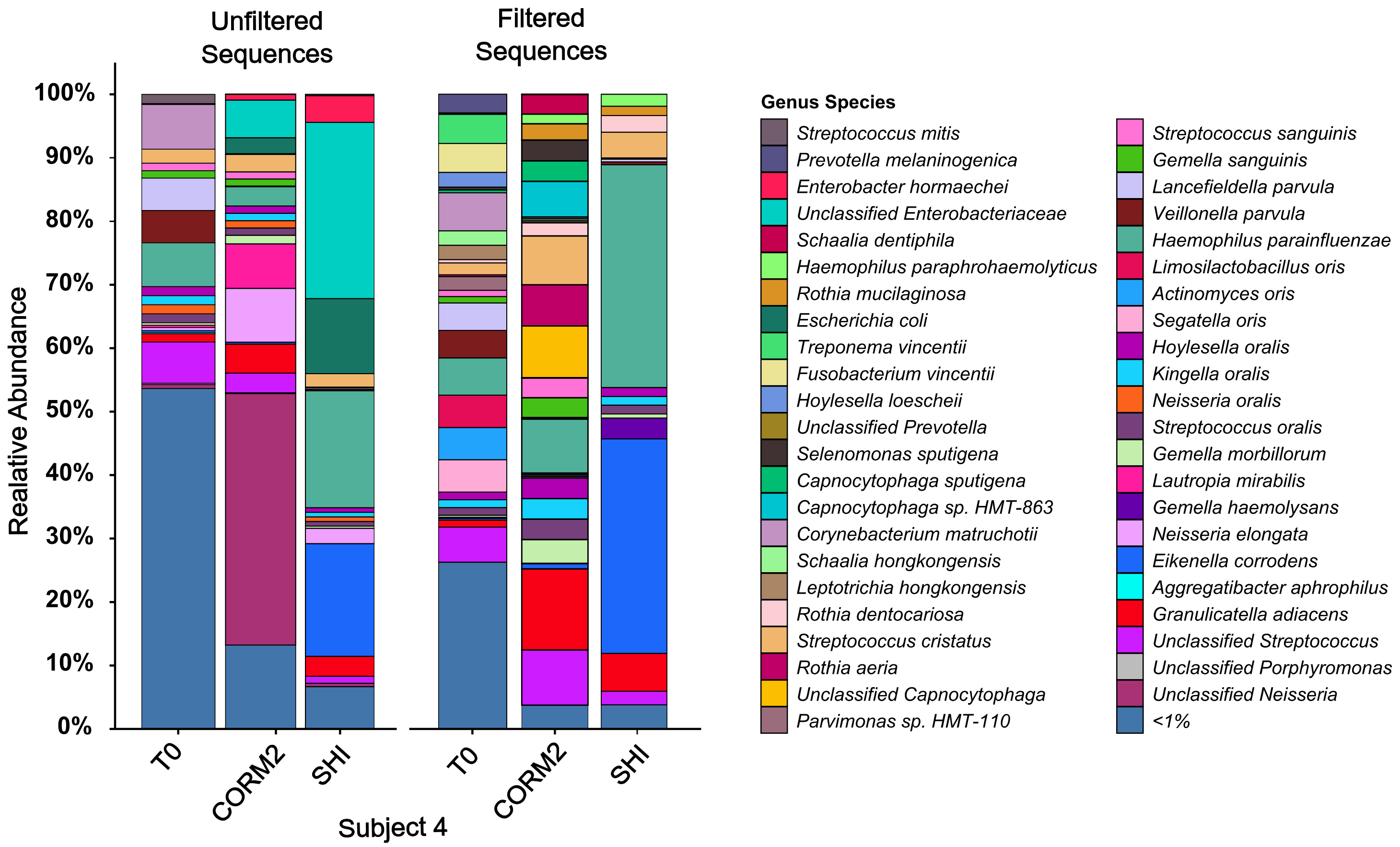


**Figure S4.** See figure 3 for associated data. Total percent abundance bar graph of identified species within samples (n=7 per bar graph) grown in CORM2 media compared to samples grown in SHI media post 3 days. “Unfiltered” refers to dataset where ASV’s belonging to genus such as Neisseria (magenta and pink), Enterobacteriaceae ( bright red and teal) and Lautropia mirabilis (hot pink) are present and compared to the same dataset where these listed ASV’s have been computationally “filtered out”. Bar graphs show underlying diversity is still present when these sequences are filtered out of the dataset.

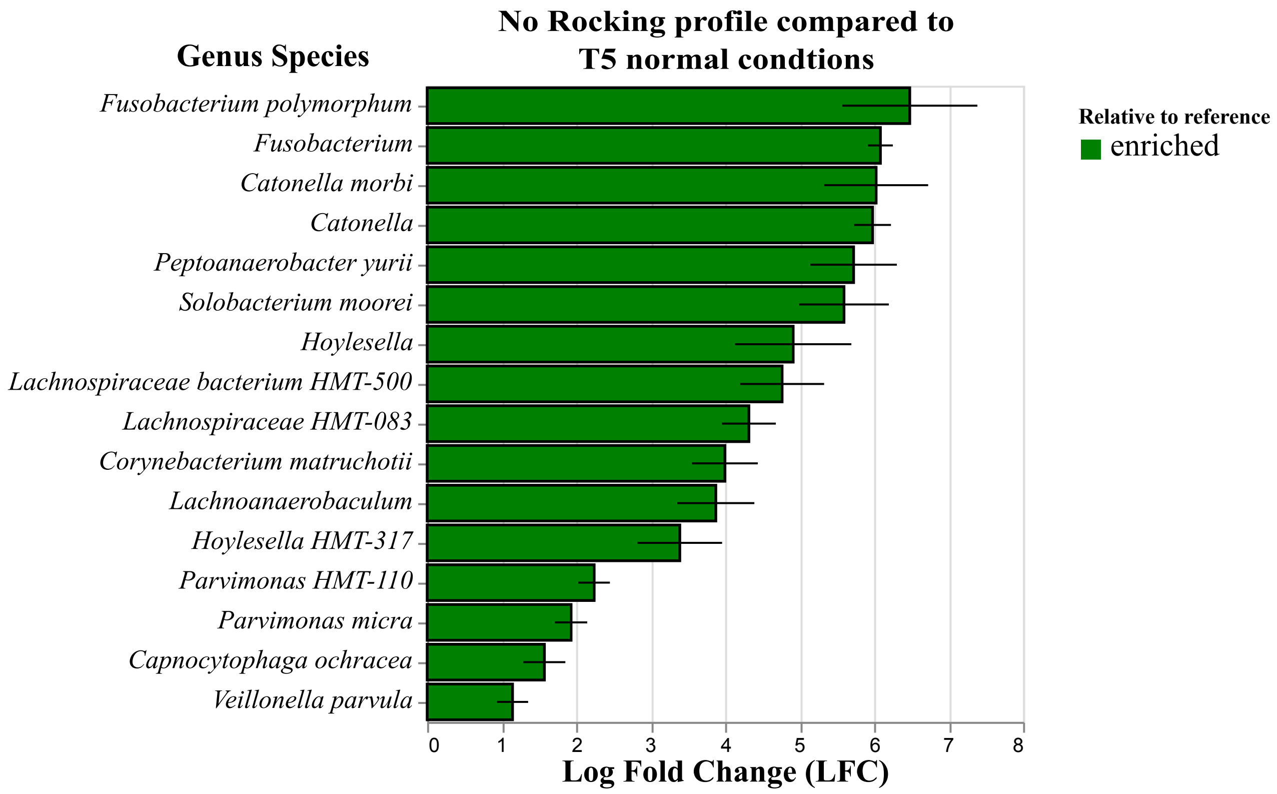


**Figure S5.** See figure 4 for associated data. Analysis of Composition of Microbiomes with Bias Correction (ANCOM-BC) between samples that were incubated statically compared to samples did were on rocking platform to allow aeration. Green bars represent abundance of taxa that are enriched in statically incubated (T5nr) samples. Red shows taxa that are depleted within t5nr samples. p threshold is ≤ 0.0001.

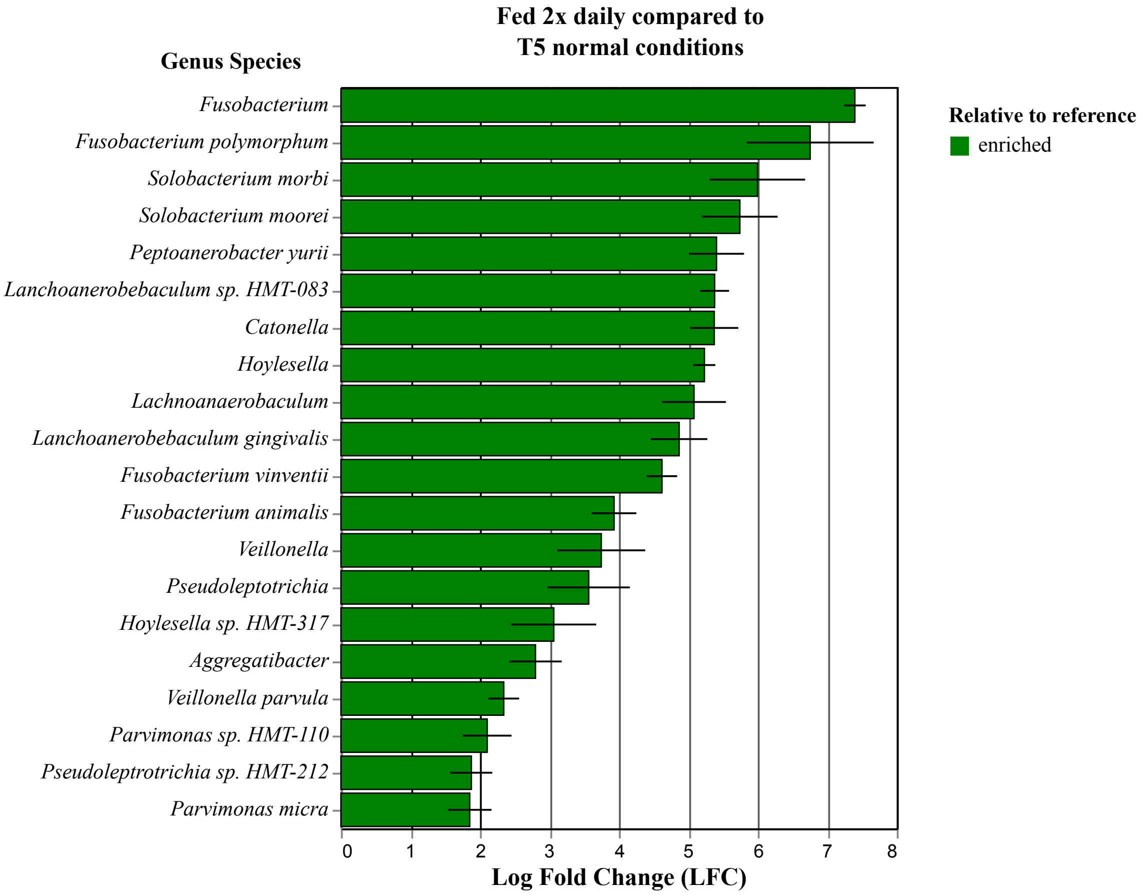


**Figure S6.** See figure 4 for associated data. Analysis of Composition of Microbiomes with Bias Correction (ANCOM-BC) between samples that were incubation entailed media replenishment occurring twice a day compared to samples where media replenishment occurred once day. Green shows taxa that are enriched in twice a day feeding (T5f2) samples. Red shows taxa that are depleted within T5f2 samples. p threshold is ≤ 0.0001.

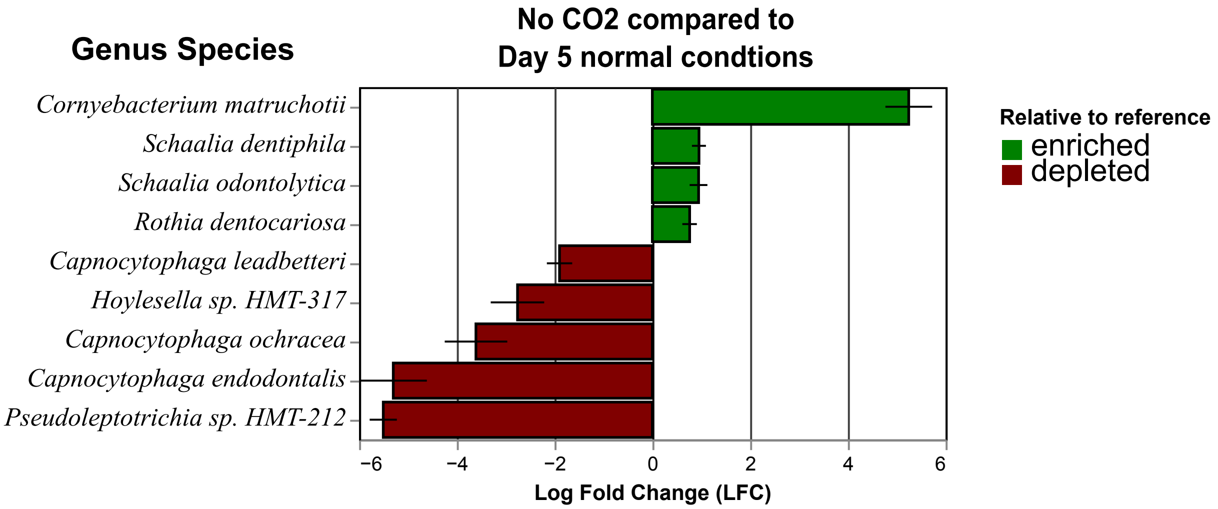


**Figure S7.** See figure 4 for associated data. Analysis of Composition of Microbiomes with Bias Correction (ANCOM-BC) between samples that were incubated without 5% $CO_2$ compared to samples that were incubated within a 5% $CO_2$ incubator. Green shows taxa that are enriched in no $CO_2$ (T5nc) samples. Red shows taxa that are depleted within T5nc. p threshold is $\leq 0.0001$.

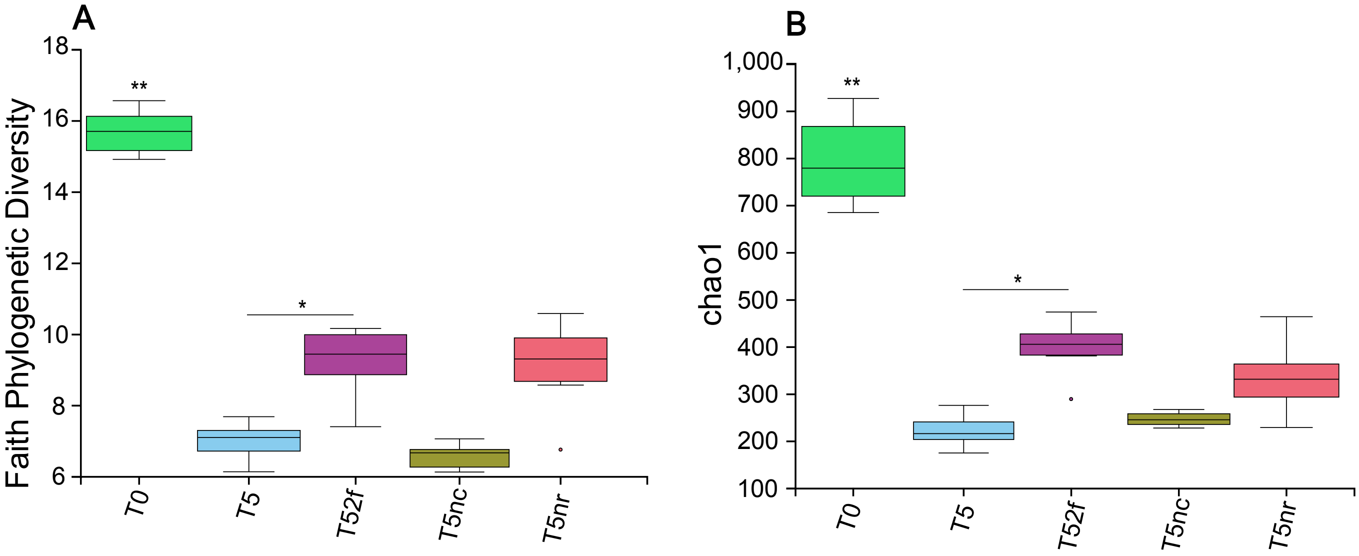


**Figure S8.** See figure 4 for associated data. **(**cont.**)** Diversity of samples grown in vacuumed filtered saliva grown in different incubation conditions, and the original T0 inoculum shown as box and whisker graphs. **T5** refers to Day 5 incubation in "normal" tested conditions listed in early sections. **T5nr** refers to samples where plaque was incubated statically noted as "not rocking". **T5nc** refers to samples which were not subjected to 5% CO2 during the 5 day incubation. **T52f** refers to samples where saliva-based media within model has been replaced 2 times a day. **(F) Faith phylogenetic distance** alpha diversity measured by total sum of branch length of all branches within the tree that spans the members found within the sample**. (H) Chao1 Distance,** measurement of total species richness based on the abundance of rare taxa within a system. Calculations were done by qiime2 via Kruskal–Wallis test (one-way ANOVA) for alpha diversity where * denotes $p \leq 0.05$, ** denotes $p \leq 0.01$, *** denotes $p \leq 0.001$ and **** denotes $p \leq 0.0001$

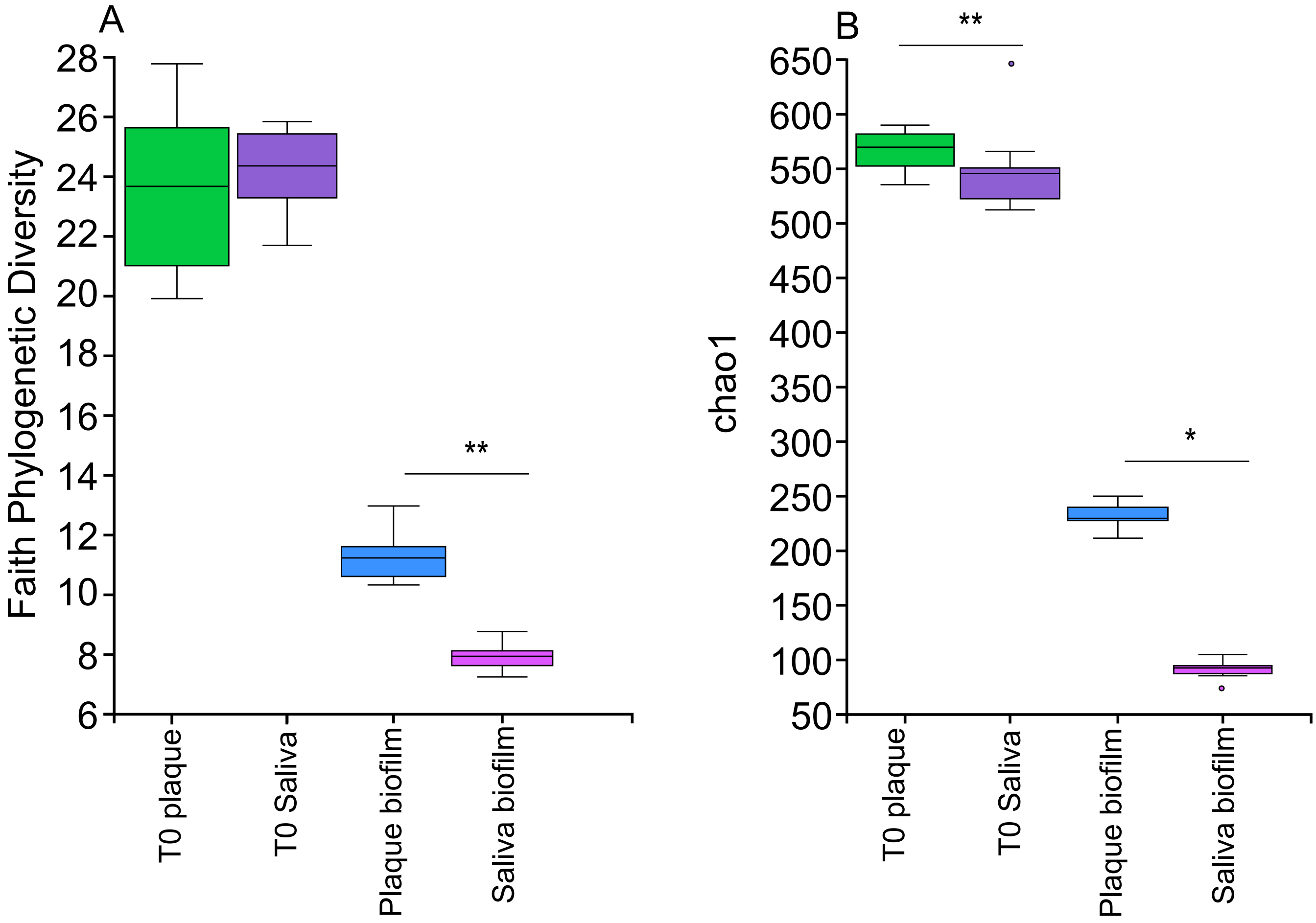


**Figure S9.** Related data in Fig. 5. (D) Faith phylogenetic distance alpha diversity measured by total sum of branch length of all branches within the tree that spans the members found within the sample. (F) Chao1 Distance, measurement of total species richness based on the abundance of rare taxa within a system. Calculations were done by qiime2 via Kruskal–Wallis test (one-way ANOVA) for alpha diversity where * denotes $p \leq 0.05$, ** denotes $p \leq 0.01$, *** denotes $p \leq 0.001$ and **** denotes $p \leq 0.0001$

### Results without *Neisseria* species computationally removed

Vacuumed sterilized saliva vs boiled saliva

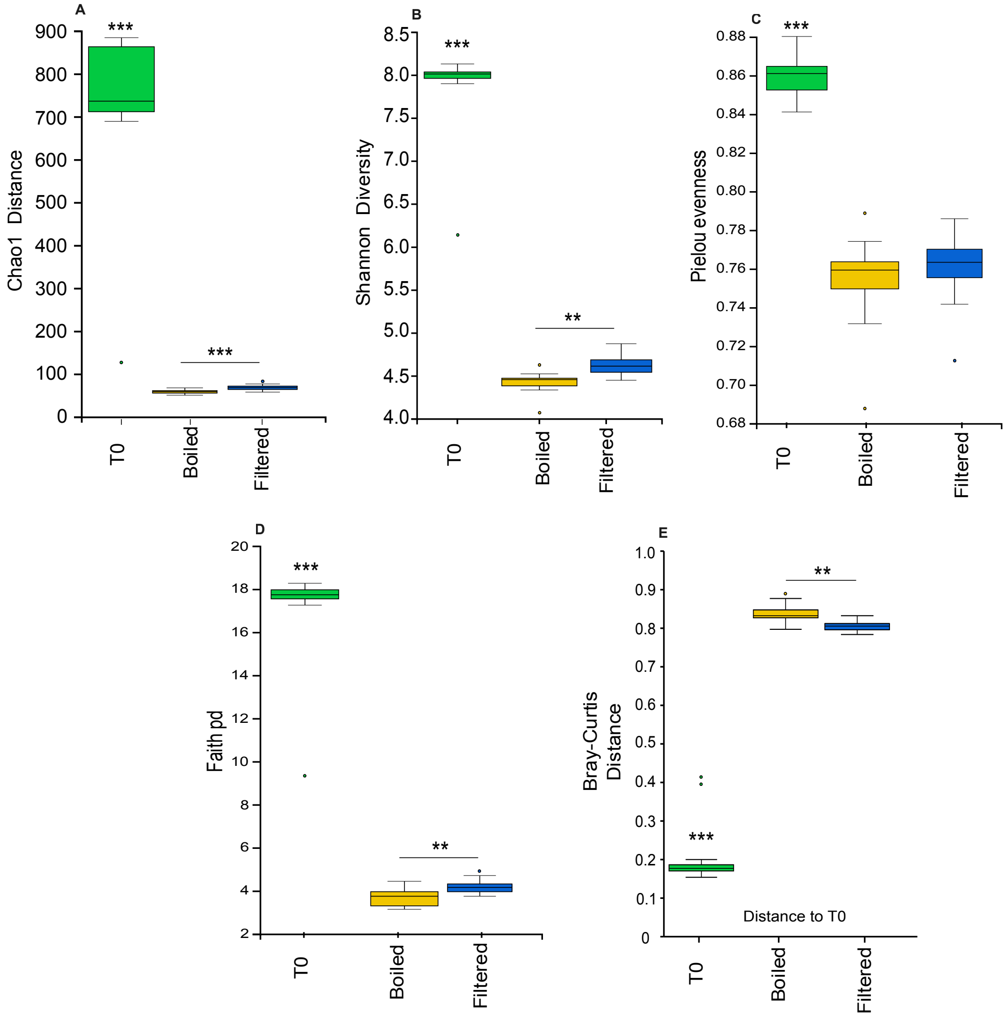


**Figure S10.** See Figure 2 for remaining dataset. (A) Chao1 Distance, measurement of total species richness based on the abundance of rare taxa within a system. (B) Shannon Chaos Index calculates the distinct species richness and evenness within a system. (C) Pielou's Evenness Distance, the measurement of how equally species within a system are distributed. (D) Faith phylogenetic distance alpha diversity measured by total sum of branch length of all branches within the tree that spans the members found within the sample. Calculations were done by qiime2 via Kruskal–Wallis test (one-way ANOVA), where * denotes $p \leq 0.05$, ** denotes $p \leq 0.01$, and *** denotes $P \leq 0.001$. (E) Bray-Curtis Dissimilarly, measurement of differences (beta diversity) of samples within a system to T0 samples. Calculation of Bray-Curtis were done in Qiime2 via Pairwise PERMANOVA test

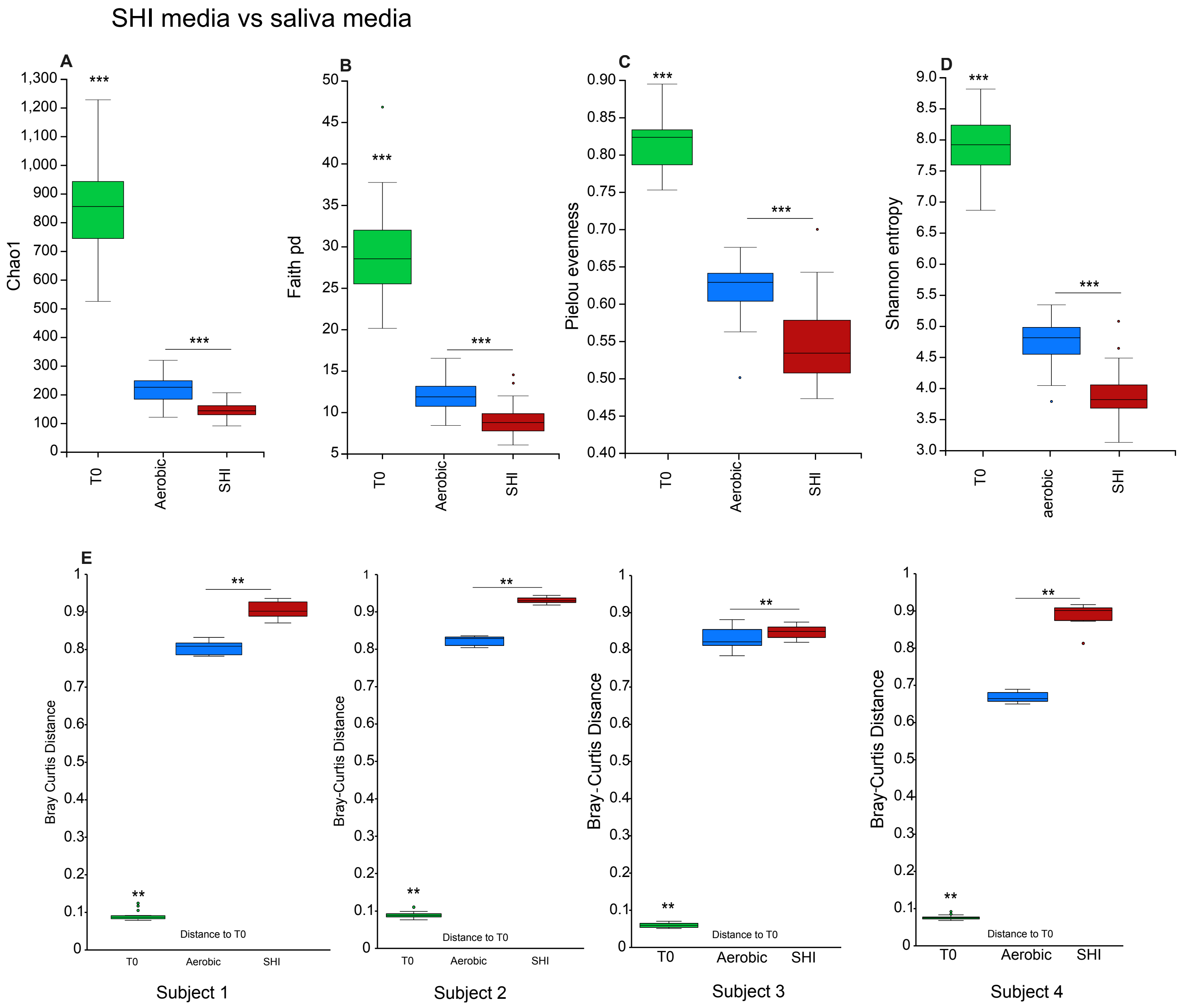


**Figure S11.** See figure 3 for associated data. (A) Chao1 Distance, measurement of total species richness based on the abundance of rare taxa within a system. (B) Faith phylogenetic distance alpha diversity measured by total sum of branch length of all branches within the tree that spans the members found within the sample (C) Pielou's Evenness Distance, the measurement of how equally species within a system are distributed. (D) Shannon Chaos Index calculates the distinct species richness and evenness within a system.  Calculations were done by qiime2 via Kruskal–Wallis test (one-way ANOVA), where * denotes $p \leq 0.05$, ** denotes $p \leq 0.01$, and *** denotes $P \leq 0.001$. (E) Bray-Curtis Dissimilarly, measurement of differences (beta diversity) of samples within a system to T0 samples. Calculation of Bray-Curtis were done in  Qiime2 via Pairwise PERMANOVA test

2x feeding/no $CO_2$/static incubation

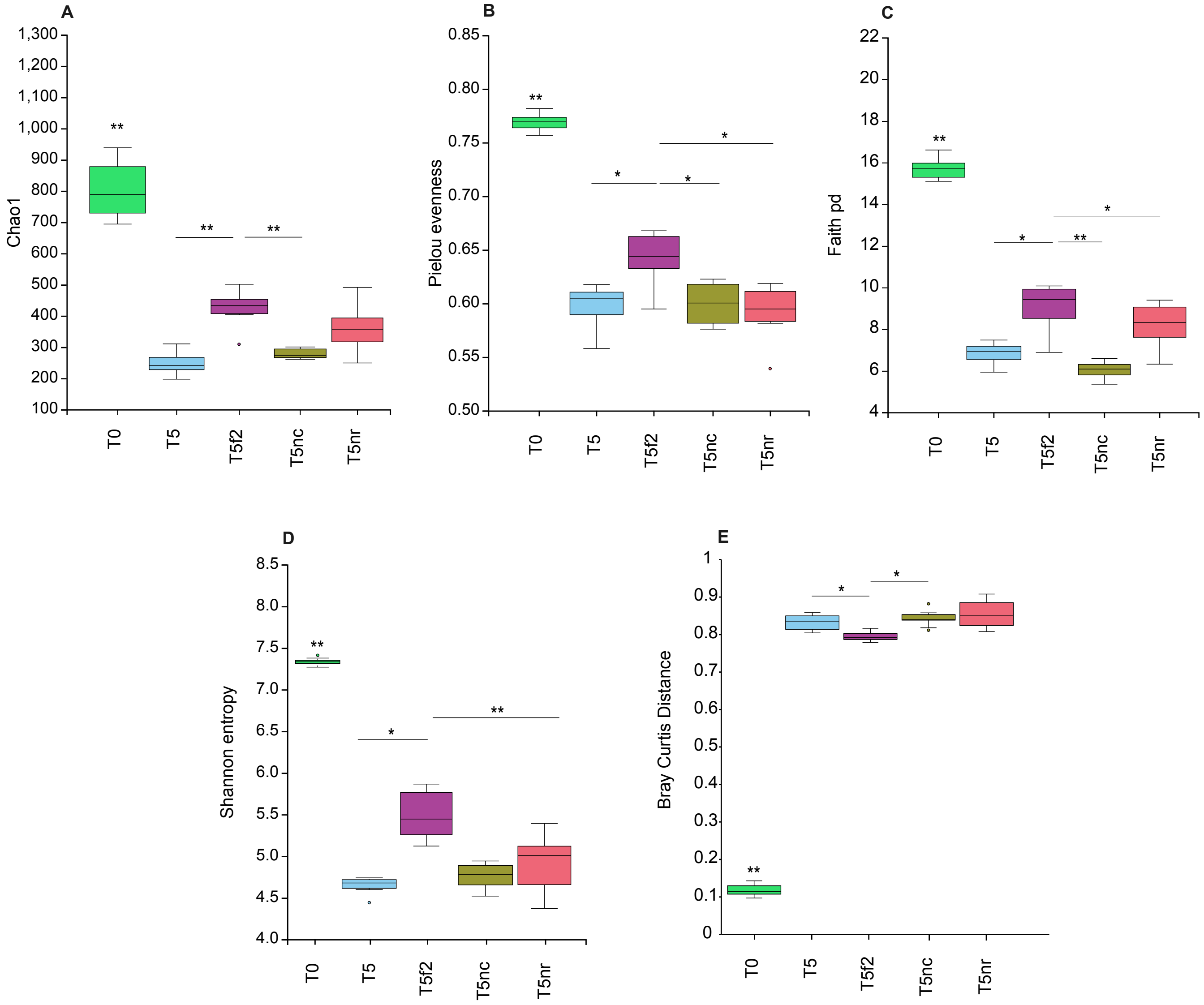


**Figure S12.** See figure 4 for associated data. (A) Chao1 Distance, measurement of total species richness based on the abundance of rare taxa within a system. (B) Pielou's Evenness Distance, the measurement of how equally species within a system are distributed. (C) Faith phylogenetic distance alpha diversity measured by total sum of branch length of all branches within the tree that spans the members found within the sample (D) Shannon Chaos Index calculates the distinct species richness and evenness within a system. Calculations were done by qiime2 via Kruskal–Wallis test (one-way ANOVA), where * denotes $p \leq 0.05$, ** denotes $p \leq 0.01$, and *** denotes $P \leq 0.001$. (E) Bray-Curtis Dissimilarly, measurement of differences (beta diversity) of samples within a system to T0 samples. Calculation of Bray-Curtis were done in Qiime2 via Pairwise PERMANOVA test.

Saliva inoculation vs Plaque inoculation

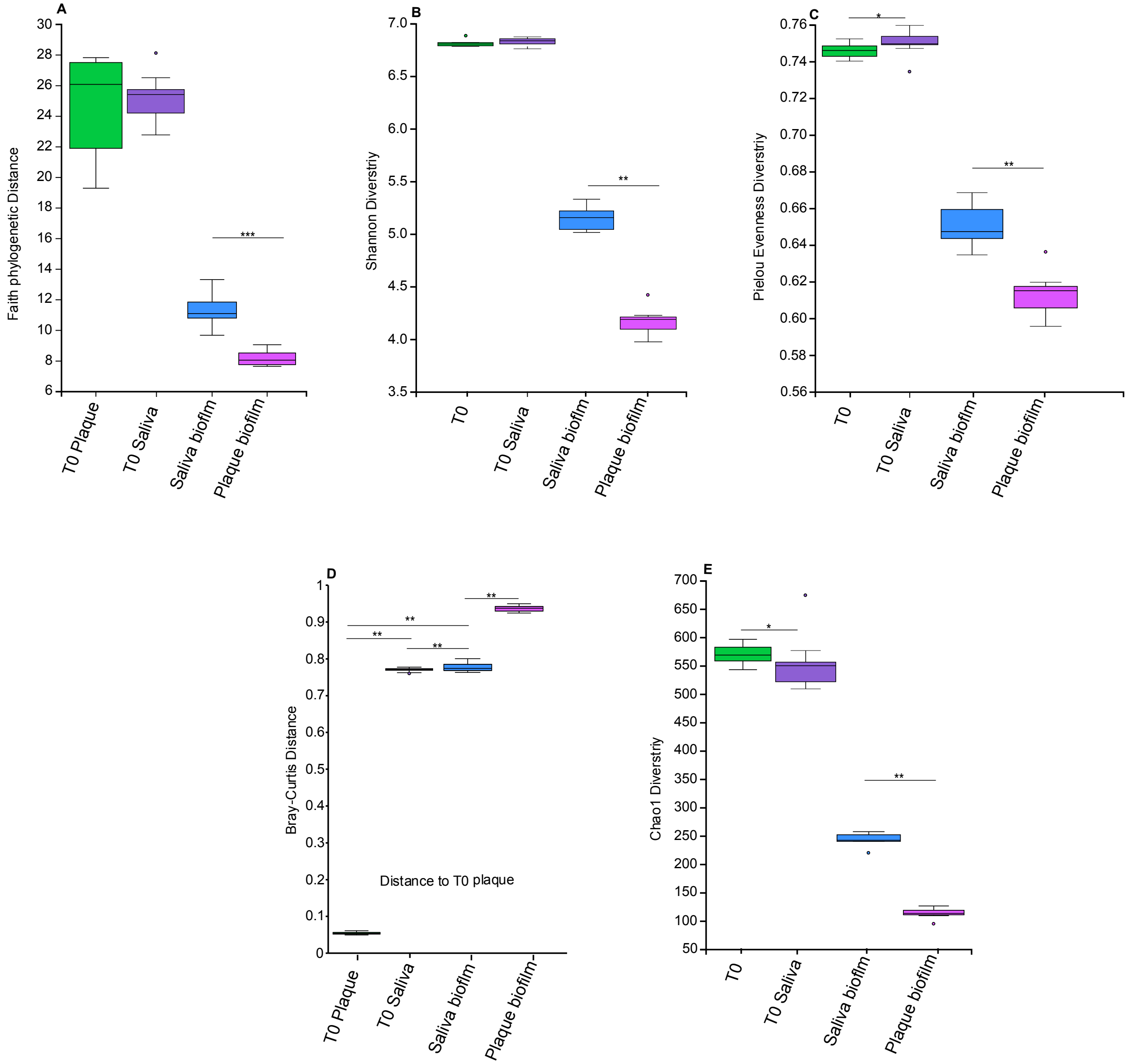


**Figure S13.** Related data in Fig. 5. Calculations were done by qiime2 via Kruskal–Wallis test (one-way ANOVA) and Bray-Curtis calculations were performed via Pairwise PERMANOVA test, where * denotes $p \leq 0.05$, ** denotes $p \leq 0.01$, and *** denotes $P \leq 0.001$. (A) Faith phylogenetic distance alpha diversity measured by total sum of branch length of all branches within the tree that spans the members found within the sample. (B) Shannon Chaos Index calculates the distinct species richness and evenness within a system (C) Pielou’s Evenness Distance, the measurement of how equally species within a system are distributed. (D) Bray-Curtis Dissimilarly, measurement of differences (beta diversity) of samples within a system to T0 samples. (E) Chao1 Distance, measurement of total species richness based on the abundance of rare taxa within a system.

### Discussion

*Sample size calculations by Power analysis*

We performed a power analysis using the difference between two independent means (standard deviation of boiled saliva vs  vacuumed filtered saliva in our case) *a priori* to gauge proper sample sizes needed for further experiments. To find the sample size which has sufficient power in we used calculated distance values from our QIIME2 analysis which was used to create the Chao1 α-diversity plots (Figure 2 supplemental), we performed a power analysis to determine Cohens δ (Cohen, 1988) in order to calculate effect and sample size:

$$\delta = \frac{\bar{x}_1 - \bar{x}_2}{\sqrt{\frac{\sigma_1^2 + \sigma_2^2}{2}}}$$

where $\bar{x}$ is the difference of mean values in observed groups and $\sigma$ is the standard deviation of the scored values. We used these values within the program G*power (Faul et al., 2007) to calculate our effect size and sample size. We initially conducted our tests based on our Chao1 calculation from this dataset using a sample size of 11 per condition. Our initial calculations indicated that 7 samples per condition would give power up to 83% with an effect size of 1.14, a large effect. However, these sample numbers are only sufficient for Chao1. Subsequent analyses showed that each diversity metric requires individual assessment (Kers & Saccenti, 2022; Rahman et al., 2023; Serdar et al., 2021). Unless stated otherwise, all further tests were completed with ≥6 samples per comparison condition, leading to possibly underpowered results for some metris. For example, *post hoc* calculation for Shannon diversity with the filtered saliva and vacuumed saliva dataset showed an effect size of 0.7, giving power of 20%, for Faith's an effect size of 1.49 with a power of 72%, Pielou evenness has an effect size of 1.25 and power of 57%. We determined that we can achieve 80% power for moderate effect sizes using samples of n=6 to n=37 depending on the metric used. It is also important to note that *post hoc* power analyses calculation can be misleading, when p-values are non-significant, *post hoc* power is low, often misleading researchers to mistake true absence of effect for insufficient study power. Thus, this model at sample sizes of ≥6 is sufficient to determine large effect sizes, but minor changes in composition should be further quantified by qPCR or other means. However, it is important to note that most current oral biofilm models cited here do not list nor mention sample sizes, power, or effect size (Cohens δ) (Bradshaw et al., 1996; Giertsen et al., 2011; B. Guggenheim et al., 2001, 2004; M. Guggenheim et al., 2001; Naginyte et al., 2019; Shapiro et al., 2002), even with human models (Utomo et al., 2024), with the exception of three papers (Edlund et al., 2013; Takeshita et al., 2015; Tu et al., 2020). Two of these papers briefly mention the importance but appear to not have shown the calculations for low effect sizes (Edlund et al., 2013; Tu et al., 2020) while the third (Takeshita et al., 2015) claims their sample size was too low for definitive statements amongst the differences of caries free vs caries positive individuals, but was suggestive enough for further experimental inquiries. Although our model has this limitation, it provides a foundation for biofilm researchers to further refine.

**Materials and Methods**

**<u>Media formulation</u>**

<u>SHI medium</u>: proteose peptone 10 g/L; trypticase peptone 5.0 g/L; yeast extract 5.0 g/L; KCl 2.5 g/L; sucrose 5 g/L; haemin 5 mg/L; VitK 1 mg/L; urea 0.06 g /L, arginine 0.174 g/L; mucin (type III, porcine, gastric) 2.5 g/L; sheep blood 5% and N -acetylmuramic acid (NAM) 10 mg/L.

<u>CORM</u>: (Teknova EZ_RICH (Teknova, USA) with saliva and other amendments): 50 mL - EZ rich defined media (Mops buffer 40 mM, Potassium phosphate dibasic 1.32 mM, 10X ACGU solution 5mL, 5X Supplement EZ solution 10mL), Lipoic acid 484 nM, Folic acid 2.26 uM, NAD 4.21 nM, Riboflavin 2.65 nM, 1000x amino acid + nucleotide block (see table below) 50 µL, Vitamin block (see table below) 50 µL. Type I water 28.25 mL.

<u>CORM2</u>**:** 40 mL – 100% pooled saliva, 10mL. 10X MOPS buffer (400mM MOPS) (Teknova, Inc), 4 mL. 1% Yeast extract (w/v), 4 mL. 100% heat inactivated human serum, 200 µL. 100 mM sucrose, 100 µL. <u>All media were if needed, adjusted to pH 7.5</u> prior to filter sterilization.

<u>Vitamin block</u>

| Vitamins/factors | 1L of 1000x, (g) | 10 ml of 1000X, (mg) |
|---|---|---|
| Choline chloride | 50.0000 | 500 |
| Beta-alanine | 10.0000 | 100 |
| Pyridoxal | 1.0000 | 10 |
| Pyridoxine HCl | 1.0000 | 10 |
| Pyridoxamine diHCl | 1.0000 | 10 |
| Spermidine triHCl | 1.0000 | 10 |
| Nicotinic acid | 1.0000 | 10 |
| Nicotinamide | 1.0000 | 10 |
| Calcium pantothenate | 1.0000 | 10 |
| Spermine tetraHCl | 1.0000 | 10 |
| Thiamine HCl | 1.0000 | 10 |
| myo-Inositol | 10.0000 | 100 |

| | | |
|---|---|---|
| Nicotinamide adenine dinucleotide | 1.0000 | 10 |
| p-Aminobenzoic acid | 0.1000 | 1 |

This solution requires high pH for components to dissolve so we added NaOH chips (~2-3) for components that did not dissolve completely while stirring at 25 °C, until fully dissolved.

Amino acid – nucleotide block

| **Amino Acids + Nucleotides**<br>not in teknova MOPS | 10mL of 1000X<br>(mg) |
|---|---|
| L-proline | 1000 |
| L-cystine | 50 |
| Thymine | 100 |
| Xanthine | 100 |

These were added in addition to amino acids and nucleotides already present in the commercial base medium formulation.

**Scripts used**

**1 - Demux**

**#imports paired end sequences (r1.qza + r2.qza) into casava file structure to use in qiime2**

```
#!/bin/bash

rawdir=/YOUR/RAW/READS/HERE

visdir=/YOUR/VISUALS/HERE


qiime tools import \
 --type 'SampleData[PairedEndSequencesWithQuality]' \
 --input-path $rawdir \
 --input-format CasavaOneEightSingleLanePerSampleDirFmt \
 --output-path $rawdir/demux-file.qza


qiime demux summarize \
 --i-data $rawdir/demux-file.qza \
 --o-visualization $visdir/demux-summary.qzv
```

**2 – Dada2 - denoise**

**#dada2 filters out noise within sequencing data, need to use the previous ".qzv" made to figure out p-trim-left and p-trunc-len**

```
#!/bin/bash

rawdir=/YOUR/RAW/READ/DIRECTORY/HERE

procdir=/YOUR/PROCESSED/DIRECTORY/HERE

visdir=/YOUR/VISUAL/DIRECTORY/HERE
```

**#Left and right forward and reverse values are gain from the demux-file from the previous script**

```
qiime dada2 denoise-paired \
  --i-demultiplexed-seqs $rawdir/demux-file.qza \
  --p-trim-left-f VALUES-HERE \
  --p-trunc-len-f  VALUES-HERE\
  --p-trim-left-r  VALUES-HERE\
  --p-trunc-len-r  VALUES-HERE\
  --o-representative-sequences $procdir/denoised-rep-seqs.qza \
  --o-table $procdir/denoised-rep-table.qza \
  --o-denoising-stats $procdir/denoise-stats.qza


qiime metadata tabulate \
  --m-input-file $procdir/denoise-stats.qza \
  --o-visualization $visdir/denoise-stats.qzv
```

**#3 - summarize script**

**#Used to gain sampling depth for rarefaction, also used to get featureID for specific sequences used later when needing to tweak classifier**

```
#!/bin/bash

#SBATCH -t 20:00:00

#SBATCH --nodes=1 --ntasks-per-node=1

#SBATCH --mem=24g

#SBATCH --mail-type=BEGIN,END,FAIL


proc=/YOUR/PROCESSED/DIRECTORY/HERE

meta=/YOUR/METADATA/FILES/HERE

visdir=/YOUR/VISUAL/DIRECTORY/HERE


qiime feature-table summarize \

  --i-table $proc/denoised-rep-table.qza \

  --m-sample-metadata-file $meta/metadata.tsv \

  --o-visualization $proc/denoised-feature-table-sum.qzv


qiime feature-table tabulate-seqs \

  --i-data $proc/denoised-rep-seqs.qza \

  --o-visualization $proc/denoised-seq-visual.qzv
```

**#4 - Alpha and Beta diversity graphs**

**#used to make box and whisker plots and PCOAs of multiple types, check qiimes2 website for more graphs not shown here**

```
#!/bin/bash

#SBATCH -t 4:00:00

#SBATCH --nodes=1 --ntasks-per-node=1

#SBATCH --mem=24g

#SBATCH --mail-type=BEGIN,END,FAIL


procdir=/YOUR/PROCESSED/DIRECTORY/HERE

metadir=/YOUR/METADATA/HERE

cmr="core-metric-results"


qiime phylogeny align-to-tree-mafft-fasttree \

  --i-sequences $procdir/denoised-rep-seqs.qza \

  --o-alignment $procdir/aligned-rep-seqs.qza \

  --o-masked-alignment $procdir/masked-aligned-rep-seqs.qza \

  --o-tree $procdir/unrooted-tree.qza \

  --o-rooted-tree $procdir/rooted-tree.qza


qiime diversity core-metrics-phylogenetic \

  --i-phylogeny $procdir/rooted-tree.qza \

  --i-table $procdir/taxa-filtered-table-FSBS.qza \

  --p-sampling-depth SAMPLE-DEPTH-FROM-PREVIOUS-SCRIPT \

  --m-metadata-file $metadir/metadata.tsv \

  --output-dir $procdir/$cmr
```

```
qiime diversity alpha \
  --i-table $procdir/denoised-feature-table.qza \
  --p-metric chao1 \
  --o-alpha-diversity $procdir/$cmr/chao1_vector.qza

qiime diversity alpha-group-significance \
  --i-alpha-diversity $procdir/$cmr/chao1_vector.qza \
  --m-metadata-file $metadir/metadata.tsv \
  --o-visualization $procdir/$cmr/chao1_group-significance.qzv

qiime diversity alpha-group-significance \
  --i-alpha-diversity $procdir/$cmr/faith_pd_vector.qza \
  --m-metadata-file $metadir/metadata.tsv \
  --o-visualization $procdir/$cmr/faith-pd-media-significance.qzv

qiime diversity alpha-group-significance \
  --i-alpha-diversity $procdir/$cmr/shannon_vector.qza \
  --m-metadata-file $metadir/metadata.tsv \
  --o-visualization $procdir/$cmr/shannon_vector.qzv

qiime diversity alpha-group-significance \
 --i-alpha-diversity $procdir/$cmr/evenness_vector.qza \
 --m-metadata-file $metadir/metadata.tsv \
 --o-visualization $procdir/$cmr/evenness-media-significance.qzv

array=( unweighted_unifrac_distance_matrix weighted_unifrac_distance_matrix
bray_curtis_distance_matrix )
```

```
for i in "${array[@]}"
do

qiime diversity beta-group-significance \
--i-distance-matrix $procdir/$cmr/$i.qza \
--m-metadata-file $metadir/metadata.tsv \
--m-metadata-column COLUMN-FROM-YOUR-METADATA \
--o-visualization $procdir/$cmr/$i.qzv \
--p-pairwise

done
```

**#5- Classifier building**

**#uses HOMD for our purpose, need specific files from HOMD site**

**#(https://www.homd.org/download/download/refseq) (such as taxa in the form of .fasta and .txt)**

**#remake your classifier when looking at data in the future as the online database updates**

**#also need your primer forward and reverse sequences used to make the classifier unique to your purpose in this case we used V1-V2 universal primers**

```
#!/bin/bash

#SBATCH -t 48:00:00

#SBATCH --nodes=1 --ntasks-per-node=1

#SBATCH --mem=288g

#SBATCH --mail-type=BEGIN,END,FAIL


classdir=/YOUR/CLASSIFIER/BUILDING/DIRECTORY/HERE

class=/YOUR/FINISHED/CLASSIFIER/DIRECTORY/HERE


if qiime tools import \

  --type 'FeatureData[Sequence]' \

  --input-path $classdir/HOMD_16S_rRNA_RefSeq_V16.01_full.fasta \

  --output-path $classdir/HOMD_16S_otu.qza ; then

  echo "fasta to qza made"

fi


if qiime tools import \

  --type 'FeatureData[Taxonomy]' \

  --input-path $classdir/HOMD_taxa.txt \

  --input-format HeaderlessTSVTaxonomyFormat \

  --output-path $classdir/ref_taxonomy.qza ; then

  echo "reference taxanonmy for classifier made"
```

```
fi

echo "files for classifier made successfully!"

if qiime rescript cull-seqs \
    --i-sequences $classdir/HOMD_16S_otu.qza \
    --o-clean-sequences $classdir/HOMD_16S_otu-cleaned.qza ; then
    echo "rescript culling done"
fi

if    qiime rescript filter-seqs-length-by-taxon \
    --i-sequences $classdir/HOMD_16S_otu-cleaned.qza \
    --i-taxonomy $classdir/ref_taxonomy.qza \
    --p-labels Archaea Bacteria Eukaryota \
    --p-min-lens 900 1200 1400 \
    --o-filtered-seqs $classdir/HOMD_16S_otu-seqs-filt.qza \
    --o-discarded-seqs $classdir/HOMD_16S_otu-seqs-discard.qza ; then
    echo "filter seq done"
fi

if qiime rescript dereplicate \
    --i-sequences $classdir/HOMD_16S_otu-seqs-filt.qza \
    --i-taxa $classdir/ref_taxonomy.qza \
    --p-mode 'uniq' \
    --o-dereplicated-sequences $classdir/HOMD_16S_otu-seqs-filt-derep-lca.qza \
    --o-dereplicated-taxa $classdir/ref_taxonomy-derep-lca.qza ;then
    echo "dereplicated completed"
fi

if  qiime feature-classifier extract-reads \
```

```
    --i-sequences $classdir/HOMD_16S_otu-seqs-filt-derep-lca.qza \
    --p-f-primer AGAGTTTGATYMTGGCTCAG \
    --p-r-primer TGCTGCCTCCCGTAGRAGT \
    --p-n-jobs 2 \
    --p-read-orientation 'forward' \
    --o-reads $classdir/HOMD-seqs-v1-v2.qza ; then
    echo "V1V2 primers aligned to classifier"
fi

if qiime rescript dereplicate \
    --i-sequences $classdir/HOMD_16S_otu-seqs-filt.qza \
    --i-taxa $classdir/ref_taxonomy.qza \
    --p-mode 'uniq' \
    --o-dereplicated-sequences $classdir/HOMD_16S_otu-seqs-filt-derep-lca.qza \
    --o-dereplicated-taxa $classdir/ref_taxonomy-derep-lca.qza ;then
    echo "dereplicated again"
fi

if qiime feature-classifier fit-classifier-naive-bayes \
    --i-reference-reads $classdir/HOMD_16S_otu-seqs-filt-derep-lca.qza \
    --i-reference-taxonomy $classdir/ref_taxonomy-derep-lca.qza \
    --o-classifier $class/HOMD-classifier-2025.qza ; then
    echo "classifier made, use this qza with your rep-seq now!"
fi
```

**#6 training your classifier on your data**

**# this makes the taxonomy.qza that combines the classifier data with your sequences, need to use specific taxonomy.qza per dataset (e.g. cannot make "one for all" type file across datasets)**

```
#!/bin/bash
#SBATCH -t 4:00:00
#SBATCH --nodes=1 --ntasks-per-node=1
#SBATCH --mem=24g
#SBATCH --mail-type=BEGIN,END,FAIL

#Must train classifier per each data set used

clsdir=/CLASSIFIER/DIRECTORY/HERE
procdir=/PROCESSED/DIRECTORY/HERE
visdir=/VISUAL/DIRECTORY/HERE

qiime feature-classifier classify-sklearn \
  --i-classifier $clsdir/HOMD-classifier-2025.qza \
  --i-reads $procdir/denoised-rep-seqs.qza \
  --o-classification $procdir/taxonomy.qza

qiime metadata tabulate \
  --m-input-file $procdir/taxonomy.qza \
  --o-visualization $procdir/taxonomy.qzv
```

**#7 taxonomic bar graph**

**#makes percent abundance graph with your taxonomy file, metadata and your denoised feature table**

```
#!/bin/bash
#SBATCH -t 4:00:00
#SBATCH --nodes=1 --ntasks-per-node=1
#SBATCH --mem=24g
#SBATCH --mail-type=BEGIN,END,FAIL

procdir=/YOUR/PROCESSED/DIRECTORY/HERE
taxa=/YOUR/TAXA/DIRECTORY/HERE
visdir=/YOUR/VISUAL/DIRECTORY/HERE
meta=/YOUR/METADATA/DIRECTORY/HERE

qiime taxa barplot \
  --i-table $procdir/denoised-feature-table.qza \
  --i-taxonomy $taxa/taxonomy.qza \
  --m-metadata-file $meta/metadata.tsv \
  --o-visualization $visdir/taxa-bar-plots.qzv
```

**# OPTIONAL SCRIPTS**
**#8-FIXING CLASSIFICATION**

**#use this to fix issues with classification**

**#turns taxonomy.qza into a TSV files**

**#match feature ID with IDs found in rep-seqs.qza from script 3 – summarize**

**#can BLAST specific sequences from rep-seq to confirm if annotation needs to be updated**

```
#!/bin/bash

#SBATCH -t 4:00:00

#SBATCH --nodes=1 --ntasks-per-node=1

#SBATCH --mem=24g

#SBATCH --mail-type=BEGIN,END,FAIL


taxa=/YOUR/TAXA/DIRECTORY/HERE



qiime tools export \

  --input-path $taxa/taxonomy.qza\

  --output-path $taxa/exported-taxonomy


#once fixed used this script below to turn the TSV back into a QZA and rerun the bar plot script with this new taxonomy.qza

qiime tools import \

  --input-path $taxa/taxonomy.tsv \

  --output-path $taxa/taxonomy.qza \

  --type 'FeatureData[Taxonomy]' \

  --input-format HeaderlessTSVTaxonomyFormat
```

**#9 ANCOM BAR GRAPHS**

**#used to produce ANCOM graphs to see differences of species between conditions**

**#can change significance threshold under --p-significance-threshold**

```
#!/bin/bash
#SBATCH -t 4:00:00
#SBATCH --nodes=1 --ntasks-per-node=1
#SBATCH --mem=24g
#SBATCH --mail-type=BEGIN,END,FAIL

procdir=/PROCESSED/DIRECTORY/HERE
meta=/METADATA/DIRECTORY/HERE
visdir=/VISUAL/DIRECTORY/HERE

qiime composition ancombc \
  --i-table $procdir/feature-table.qza \
  --m-metadata-file $meta/metadata.tsv \
  --p-formula COLUMN-NAME \
  --p-reference-levels COLUMN-NAME::THING-YOU-WANT-TO-COMPARE-AS-REFERNECE \
  --o-differentials $procdir/ancombc.qza

qiime composition da-barplot \
  --i-data $procdir/ancombc.qza \
  --p-significance-threshold 0.0001 \
  --o-visualization $visdir/da-barplot-0001.qzv

qiime taxa collapse \
  --i-table $procdir/feature-table.qza \
```

```
  --i-taxonomy $procdir/taxonomy.qza \
  --p-level 7 \
  --o-collapsed-table $procdir/table-l7.qza

qiime composition ancombc \
  --i-table $procdir/table-l7.qza \
  --m-metadata-file $meta/metadata.tsv \
  --p-formula COLUMN-NAME \
  --p-reference-levels COLUMN-NAME::THING-YOU-WANT-TO-COMPARE-AS-
REFERENCE \
  --o-differentials $procdir/l7-ancombc.qza

qiime composition da-barplot \
  --i-data $procdir/l7-ancombc.qza \
  --p-significance-threshold 0.0001 \
  --p-level-delimiter ";" \
  --o-visualization $visdir/l7-da-barplot-0001.qzv
```

**#10 filters**

**#Filtered-scripts**

**#example of a script used to filtered metadata to make a feature table that can be used for specific samples/subjects based on users need**

**#also make summary table for use for sampling depth**

```
#!/bin/bash

#SBATCH -t 4:00:00

#SBATCH --nodes=1 --ntasks-per-node=1

#SBATCH --mem=24g

#SBATCH --mail-type=BEGIN,END,FAIL


procdir=/YOUR/PROCESSED/FILES/HERE

meta=/YOUR/METADATA/FILES/HERE

filter=/YOUR/FILTERED/DATA/HERE


qiime feature-table filter-samples \
  --i-table $procdir/rep-table.qza \
  --m-metadata-file $meta/metadata.tsv \
  --p-where "[COLUMN-NAME] IN ('SAMPLE','SAMPLE-2','SAMPLE-3','SAMPLE-4')" \
  --o-filtered-table $filter/filtered-meta-table.qza


qiime feature-table summarize \
  --i-table $filter/filtered-meta-table.qza \
  --m-sample-metadata-file $meta/metadata.tsv \
  --o-visualization $filter/filtered-meta-table-summary.qzv
```

**Work Cited**

Bradshaw, D. J., Marsh, P. D., Schilling, K. M., & Cummins, D. (1996). A modified chemostat system to study the ecology of oral biofilms. *Journal of Applied Bacteriology*, *80*(2), 124–130. https://doi.org/10.1111/j.1365-2672.1996.tb03199.x

Cohen, J. (1988). *Statistical power analysis for the behavioral sciences* (2. ed., reprint). Psychology Press.

Edlund, A., Yang, Y., Hall, A. P., Guo, L., Lux, R., He, X., Nelson, K. E., Nealson, K. H., Yooseph, S., Shi, W., & McLean, J. S. (2013). An in vitrobiofilm model system maintaining a highly reproducible species and metabolic diversity approaching that of the human oral microbiome. *Microbiome*, *1*(1), 25. https://doi.org/10.1186/2049-2618-1-25

Faul, F., Erdfelder, E., Lang, A.-G., & Buchner, A. (2007). G*Power 3: A flexible statistical power analysis program for the social, behavioral, and biomedical sciences. *Behavior Research Methods*, *39*(2), 175–191. https://doi.org/10.3758/bf03193146

Giertsen, E., Arthur, R. A., & Guggenheim, B. (2011). Effects of Xylitol on Survival of Mutans Streptococci in Mixed-Six-Species in vitro Biofilms Modelling Supragingival Plaque. *Caries Research*, *45*(1), 31–39. https://doi.org/10.1159/000322646

Guggenheim, B., Giertsen, E., Schüpbach, P., & Shapiro, S. (2001). Validation of an in vitro Biofilm Model of Supragingival Plaque. *Journal of Dental Research*, *80*(1), 363–370. https://doi.org/10.1177/00220345010800011201

Guggenheim, B., Guggenheim, M., Gmür, R., Giertsen, E., & Thurnheer, T. (2004). Application of the Zürich Biofilm Model to Problems of Cariology. *Caries Research*, *38*(3), 212–222. https://doi.org/10.1159/000077757

Guggenheim, M., Shapiro, S., Gmür, R., & Guggenheim, B. (2001). Spatial Arrangements and Associative Behavior of Species  in an In Vitro Oral Biofilm Model. *Applied and Environmental Microbiology*, *67*(3), 1343–1350. https://doi.org/10.1128/AEM.67.3.1343-1350.2001

Kers, J. G., & Saccenti, E. (2022). The Power of Microbiome Studies: Some Considerations on Which Alpha and Beta Metrics to Use and How to Report Results. *Frontiers in Microbiology*, *12*. https://doi.org/10.3389/fmicb.2021.796025

Naginyte, M., Do, T., Meade, J., Devine, D. A., & Marsh, P. D. (2019). Enrichment of periodontal pathogens from the biofilms of healthy adults. *Scientific Reports*, *9*(1), 5491. https://doi.org/10.1038/s41598-019-41882-y

Rahman, G., McDonald, D., Gonzalez, A., Vázquez-Baeza, Y., Jiang, L., Casals-Pascual, C., Hakim, D., Dilmore, A. H., Nowinski, B., Peddada, S., & Knight, R. (2023). Determination of Effect Sizes for Power Analysis for Microbiome Studies Using Large Microbiome Databases. *Genes*, *14*(6), 1239. https://doi.org/10.3390/genes14061239

Serdar, C. C., Cihan, M., Yücel, D., & Serdar, M. A. (2021). Sample size, power and effect size revisited: Simplified and practical approaches in pre-clinical, clinical and laboratory studies. *Biochemia Medica*, *31*(1), 010502. https://doi.org/10.11613/BM.2021.010502

Shapiro, S., Giertsen, E., & Guggenheim, B. (2002). An in vitro Oral Biofilm Model for Comparing the Efficacy of Antimicrobial Mouthrinses. *Caries Research*, *36*(2), 93–100. https://doi.org/10.1159/000057866

Takeshita, T., Yasui, M., Shibata, Y., Furuta, M., Saeki, Y., Eshima, N., & Yamashita, Y. (2015). Dental plaque development on a hydroxyapatite disk in young adults observed by using a barcoded pyrosequencing approach. *Scientific Reports*, *5*(1), Article 1. https://doi.org/10.1038/srep08136

Tu, Y., Wang, Y., Su, L., Shao, B., Duan, Z., & Deng, S. (2020). In vivo Microbial Diversity Analysis on Different Surfaces of Dental Restorative Materials via 16S rDNA Sequencing. *Medical Science Monitor : International Medical Journal of Experimental and Clinical Research*, *26*, e923509-1-e923509-11. https://doi.org/10.12659/MSM.923509

Utomo, R. N. C., Palkowitz, A. L., Gan, L., Rudzinski, A., Franzen, J., Ballerstedt, H., Zimmermann, M., Blank, L. M., Fischer, H., Wolfart, S., & Tuna, T. (2024). In vitro plaque formation model to unravel biofilm formation dynamics on implant abutment surfaces. *Journal of Oral Microbiology*, *16*(1), 2424227. https://doi.org/10.1080/20002297.2024.2424227
